# BatchRefiner: fast, significant improvement in batch integration of single-cell embeddings with ensemble refinement

**DOI:** 10.64898/2026.08.21.746347

**Authors:** Daniel E. Schäffer, Helen Kang, Ekin Deniz Aksu, Daniel Edelman, Bonnie Berger

## Abstract

Data from single-cell RNA sequencing (scRNA-seq) and the Assay for Transposase-Accessible Chromatin (scATAC-seq) are high-dimensional, sparse, and undesirably capture technical variability between experiments or batches. Many analysis methods thus seek to produce a low-dimensional cell-by-feature embedding space that groups together biologically similar cells across batches while distancing dissimilar cells. Here, we introduce *ensemble refinement* for scRNA-seq and scATAC-seq embeddings, inspired by ensemble methods from statistical machine learning, and implement BatchRefiner, a fast post-processing tool to enhance batch integration. We extensively benchmark widely-used scRNA-seq embedding methods on both batch integration and biological conservation over a wide range of datasets, before and after the addition of BatchRefiner. We extend these benchmarking approaches to provide the first comprehensive benchmark of batch integration for scATAC-seq embedding methods, including BatchRefiner. Importantly, we formalize a significance statistic, which we use to demonstrate BatchRefiner’s significant improvement in batch integration across a wide range of embedding methods, atlas-scale datasets, and established metrics.

## Introduction

Single-cell RNA-sequencing (scRNA-seq) assays measure gene expression in populations of individual cells and enable biological discovery at unprecedented speed and scale [1–3]. The amount of scRNA-seq and other single-cell omics data has grown rapidly as technologies have matured and proliferated [4, 5], with scRNA-seq currently the most popular and plentiful modality. Building on high-throughput scRNA-seq efforts, single-cell atlases bring together data from large numbers of cells collected from multiple experiments, assays, and biological sources [5, 6]. The complexity in these data present not only as biological variation, but as technical variation, also known as batch effects [7, 8]. Differences in sample collection, laboratory conditions, library preparation, and sequencing technology leave unique footprints in measured gene expression profiles. This predicament has motivated the development of a multitude of single-cell batch integration methods [9–12] and also approaches to benchmark the performance of those methods [7, 8, 13–16].

Single-cell batch-integration methods commonly map gene expression profiles from different batches to a joint embedding space. Ideally, this embedding space: 1) is devoid of batch effects—cells from the same batch do not have a tendency to group together (“batch integration”) *and* 2) preserves true biological signal—biologically similar cells group together regardless of their original batch (“bio conservation”). Many metrics of bio conservation rely on cell-type labels of reference datasets, and bio conservation is therefore partially synonymous with preservation of distinct cell-type identities [8]. Current batch-integration methods achieve a range of tradeoffs between batch integration and bio conservation, and improvements in one likely come at the expense of the other [8]. Recent scRNA-seq benchmarking has made clear that batch integration, also referred to as batch correction, remains an important barrier to atlas-scale dataset integration [16, 17]. Furthermore, although users might require different levels of batch integration for different datasets and downstream applications, current batch-integration tools cannot directly adjust their level of batch correction [17, 18].

While scRNA-seq takes a collection of sequencing reads and produces a cell-by-gene matrix that counts the number of reads from each cell that map to each gene [1–3], the Assay for Transposase-Accessible Chromatin with Sequencing (ATAC-seq) is an orthogonal sequencing-based assay that can be applied to characterize genomic states at the tissue or single-cell level. With ATAC-seq, the output sequencing reads are derived from a segment of DNA that is accessible to protein binding at the time of the assay, specifically binding by the Tn5 transposase [19, 20]. ATAC-seq reads are preferentially in the vicinity of transcribed genes, especially transcriptional start sites and promoters, as well as accessible, non-transcribed regions of the genome that are generally taken to be active regulatory elements (enhancers). The results of scATAC-seq can be summarized as a cell-by-peak matrix that counts the number of reads from each cell that map to each accessible site, defined as a small positional interval in base pair coordinates with respect to a reference genome [21, 22]. Thus, the fundamental problem of converting a high-dimensional and sparse per-cell readout to a low-dimensional and *de-noised* representation is shared between scRNA-seq, scATAC-seq, and other related omics assays.

Here, we introduce *ensemble refinement* for omics embeddings through **BatchRefiner**, a post-processing approach derived from ensemble learning. We apply BatchRefiner to the embeddings output by existing methods, both for scRNA-seq and scATAC-seq, and show it significantly improves batch integration of single-cell embeddings. To robustly test the relative performance of embedding methods, we first expand current benchmarking approaches for scRNA-seq embeddings to additional widely-used methods (*e.g.*, Seurat [23]) and augment the set of representative benchmarking datasets. To explore the generalizability of BatchRefiner, we do the same for the scATAC-seq modality, which has been the subject of comparatively sparse benchmarking for batch integration [24]. We assemble a panel of nine scATAC-seq datasets from separate studies and extend the end-to-end scRNA-seq benchmarking pipeline [8] to enable evaluations of embedding methods for other modalities using shared metrics. We find that scATAC-seq specific embedding methods alone offer poor intrinsic batch correction. To address this limitation, authors have combined their embeddings with the scRNA-seq method Harmony [25] to achieve quantitatively effective batch integration, which can be statistically improved by passing it as input to BatchRefiner for post-processing. In contrast to scRNA-seq, we observe substantial variation in bio conservation even among uncorrected scATAC-seq methods, highlighting the relative complexity of and room for improvement in embedding scATAC-seq data.

The key conceptual advance of ensemble refinement is to treat dimensions of an embedding as separate “weak” embeddings, extending the notion of weak learners from ensemble methods [26, 27]. We draw inspiration from *stacked generalization* approaches [28, 29] to design BatchRefiner for *post hoc* evaluation and re-weighting of an ensemble. Our method quantifies the degree of batch effects in each embedding dimension, which is used to inform a refinement of an initial embedding to improve batch integration (Figure 1). BatchRefiner *significantly* improves batch integration for widely-used scRNA-seq and scATAC-seq integration methods, as shown by a newly-formalized significance statistic for which we leverage the statistical power of independent benchmarking datasets. We highlight that because both our comprehensive benchmarking and BatchRefiner method take a cell embedding matrix as input, they are generalizable to any current or future scRNA-seq or scATAC-seq embedding methods. As multimodal single-cell assay technologies contribute orthogonal biological information [30–32] and multiomic data reach atlas scale [5], BatchRefiner will be further generalizable to additional omics modalities and related batch-correction methods.

**Fig. 1.**
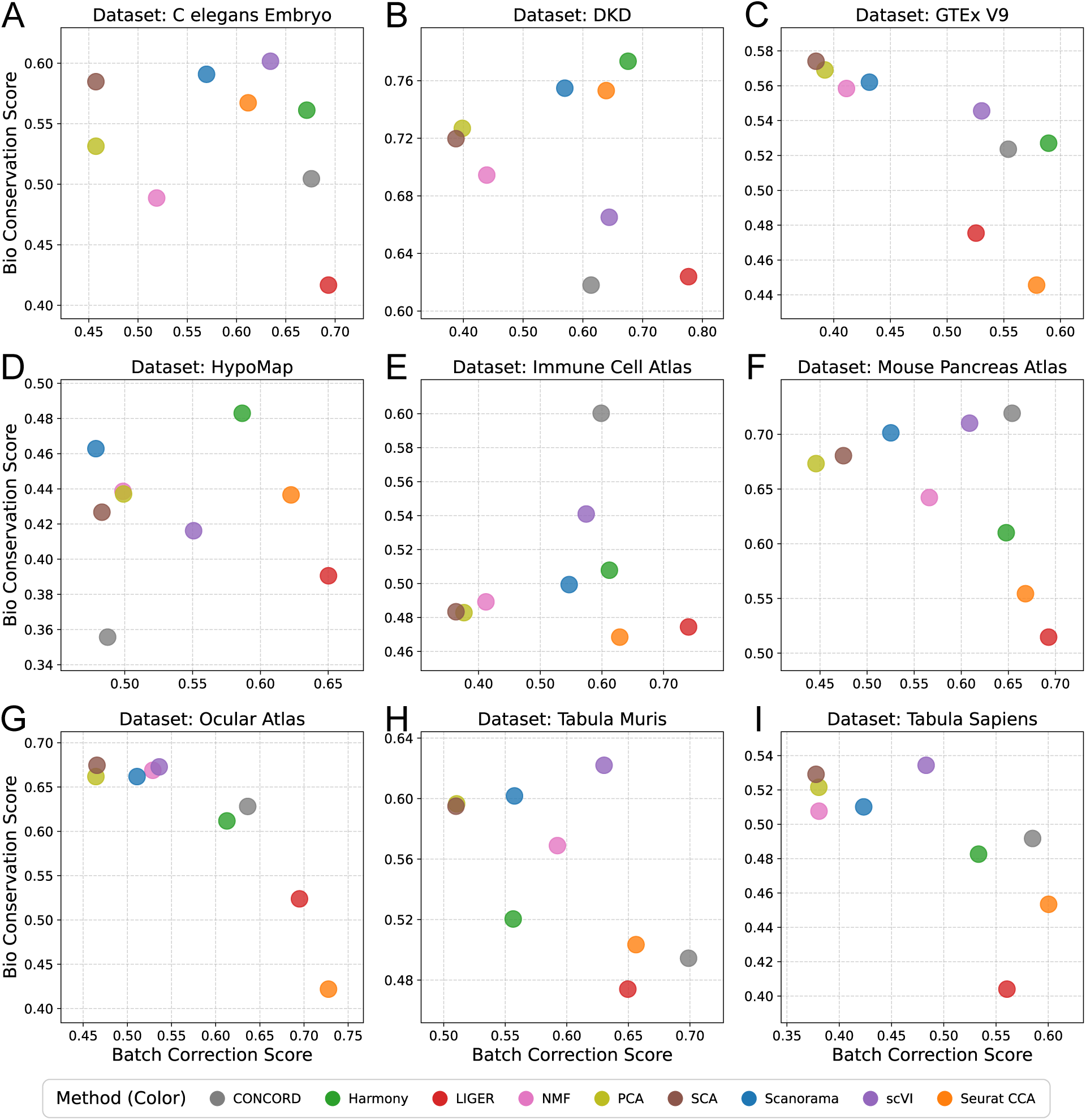
Benchmarks of scRNA-seq embedding methods on each of nine datasets. Benchmarking of embedding methods, shown as bio-conservation (y-axis) and batch-integration (x-axis, Batch Correction) scores for each of nine scRNA-seq datasets individually. These scores are the average of seven and five, respectively, individual metrics that are each scaled per-dataset (Methods). Benchmark values range from 0 to 1, with 1 being best.

## Results

### Initial Benchmarking of scRNA-seq embedding methods

To perform a comprehensive, objective benchmarking of embedding methods, we selected the state-of-the-art OpenProblems framework [16], specifically the Batch Integration Task version 2 (Methods). We extended this benchmarking approach in several ways, including to introduce new embedding methods and datasets, and to newly apply tests of statistical significance for improvement in batch correction (Methods) for selected BatchRefiner modes relative to baseline methods. By starting with an existing benchmarking platform (datasets and metrics), we aimed to obtain an objective measure of the relative performance of BatchRefiner and baseline methods.

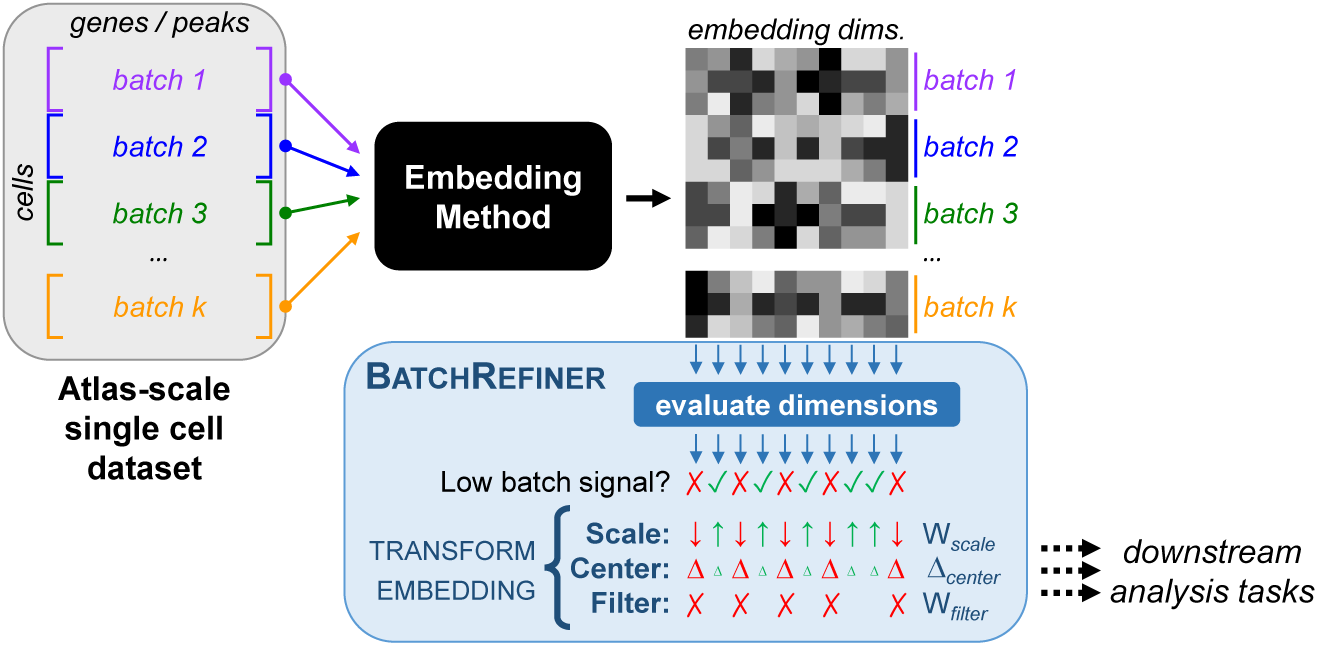
Overview of BatchRefiner: Embedding methods for scRNA-seq data (black box) take gene expression data and output a lower-dimensional embedding for each cell (top right, **A**). BatchRefiner (blue box) then performs ensemble refinement by down-scaling (down arrows representing low values in *W_scale_*, Equation 1 in Box 1), centering (large *Δ*’s representing large values in *Δ_center_*, Equation 2), or filtering out (red Xs representing 0 values in *W_filter_*, Equation 3) the dimensions with a high batch signal. This results in a refined embedding useful for downstream tasks sensitive to batch effects.

For scRNA-seq, we selected nine embedding methods as baseline approaches. We included five widely-used batch-correction methods: Harmony [25], Scanorama [33, 34], Seurat Canonical Correlation Analysis [35, 23], scVI [36, 37], and LIGER [38, 39]; as well as a recent contrastive-learning approach, CONCORD [40]. On the public framework results, scVI, Harmony, and LIGER were reported to be the three highest-scoring embedding methods [16], and along with Scanorama and Seurat score highly in previous benchmarking studies [8, 14, 15]. Of the included methods, scVI and CONCORD train dataset-specific machine learning models to embed each dataset with objectives that incorporate batch mixing. We also included three methods that do not explicitly batch correct: principal component analysis (PCA) [41], a venerable but not task-specific dimensionality reduction tool, which also serves as a pre-processing step for Harmony and Seurat; Surprisal Component Analysis (SCA) [42], and non-negative matrix factorization (NMF) [43]. We note that, of these methods, Seurat, CONCORD, SCA, and NMF are new to the OpenProblems framework, along with Scanorama’s embedding mode (Methods). We excluded label-transfer methods that make use of cell type labels, such as scANVI [44]. Such methods have an unfair advantage over other methods on datasets with accurate cell-type labels [8], and inherently could not be used to *de novo* embed newly-generated datasets.

Each embedding (a method, dataset pair) is scored on seven bio-conservation and five batch-correction metrics. Each metric produces scores between 0 and 1, with 1 being best. However, differences in dataset and metric construction can reduce the empirical ranges, so we re-scaled each metric such that 0 and 1 correspond to dataset-specific minimum and maximum scores achieved by a panel of control (randomized and perfect-information) methods (Methods, Supplementary Data). Averaged across nine scRNA-seq atlas datasets (six from OpenProblems, three added for this study; see Methods), we observe that the six baseline batch-correction methods fall generally along a frontier of performance (Table 1), which has been described previously [8]. Those methods show varying tradeoffs between bio conservation and batch integration; on average, scVI and Scanorama demonstrate the best bio conservation, whereas Seurat and LIGER provide some additional batch integration at the expense of bio conservation. However, we observe that the relative performance of these methods varies substantially between datasets (Figure 1), highlighting the importance of a varied benchmarking set and suggesting the potential importance of dataset-tuned approaches [18]. The other three methods, PCA, SCA, and NMF, demonstrate competitive bio-conservation performance but, unsurprisingly, worse batch integration (Table 1).

**Table 1.** Benchmarking of scRNA-seq embedding methods on nine datasets.

| Method | Batch-Integration Score | Bio-Conservation Score |
| --- | --- | --- |
| Concord | 0.612 | 0.548 |
| Harmony | 0.609 | 0.564 |
| LIGER | 0.665 | 0.478 |
| NMF | 0.483 | 0.562 |
| PCA | 0.436 | 0.578 |
| SCA | 0.434 | 0.585 |
| Scanorama | 0.513 | 0.594 |
| scVI | 0.577 | 0.590 |
| Seurat CCA | 0.637 | 0.512 |

### BatchRefiner: *ensemble refinement* for increased batch integration

#### Overview of BatchRefiner

We develop and evaluate three BatchRefiner modes that take distinct approaches for embedding refinement. (Figure 1, Box 1, Supplementary Table S1). First, “Scaling” (Equation 1, Box 1) tunes the relative importance of dimensions by linear scaling. Second, “Centering” (Equation 2, Box 1) subtracts from each dimension a fraction of the batch mean equal to the scaling weight for each dimension, which better preserves non-batch signal even in high-batch dimensions. Third, “Filtering” (Equation 3, Box 1) drops batch-correlated dimensions from the embedding, thus reducing the embedding dimensionality; we evaluate several thresholds for filtering, which can also be viewed as scaling with binarized (rather than linear) weights.

To assess each dimension of a given embedding, we utilize established metrics of batch integration (Methods): variance explained (*R*^2^) by the batch labels (closely related to the PCR Comparison metric [13]), or integration Local Inverse Simpson’s Index (iLISI), which measures local integration in small cell “neighborhoods” [25]. These do not require cell type labels, unlike other batch-integration metrics. Newly-generated data may not have pre-labeled cell types or may suffer from imperfect labeling, especially of rare or novel cell types. Thus, even though many benchmarking metrics rely on ground-truth cell type labels as an important source of information, it is preferable for an embedding method not to require them to embed new datasets.

#### BatchRefiner Scaling

We see consistent overall improvements from BatchRefiner scaling using batch *R*^2^ (Fisher’s combined *p* = 4.1 × 10^−29^ considering all nine methods), with significant increases in batch correction for seven of nine individual baseline methods (Bonferroni *p <* 0.05, Supplementary Table 2). Average batch-integration scores increase by up to 0.09, with small (0 − 0.04) tradeoffs in bio conservation (Figure 2A-B). For example, scaling Harmony gives it state-of-the-art batch-integration performance and places it clearly over the observed performance frontier. Scaling Scanorama, which was previously below (left) of the frontier, brings it in line with the top baseline methods (the frontier); it already scored highly in bio conservation. Seurat, and scVI show smaller, but still significant boosts, while LIGER and CONCORD have minimal change with scaling. The non-batch-correcting methods, PCA, SCA, and NMF, also benefit from scaling, although their batch-integration performance remains worse than the top standalone methods. However, scaling still increases batch integration overall even when restricted to only the six batch-correction methods (Fisher’s combined *p* = 6.1 × 10^−19^).

##### Box 1

**BatchRefiner Equations**

Let *E* ∈ ℝ^|cells|×*d*^ be embeddings, and ***b*** ∈ ℝ^*d*^ be scores quantifying batch signal for each dimension.

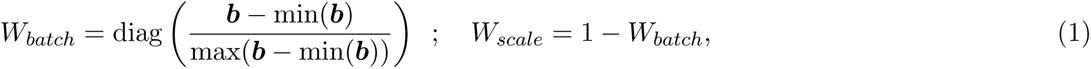

where diag(***v***) = *D* ∈ ℝ^|***v***|×|***v***|^ s.t. *D_ii_* = *v_i_*; *D_ij_* = 0 ∀*i* ≠ *j*

BatchRefiner(*E,* mode=‘scale’) = *E* × *W_scale_*

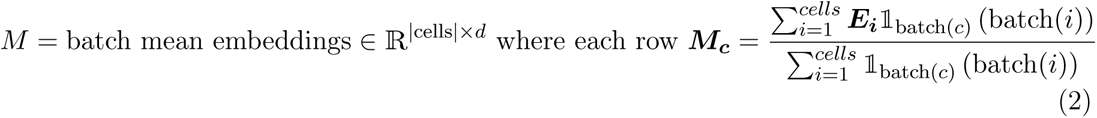

where batch(*c*) is the batch label of cell *c*

BatchRefiner(*E,* mode=‘center’) = *E* − *Δ_center_,* where *Δ_center_* = *M* × *W_batch_*

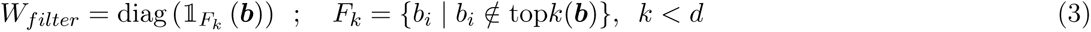

where top*k*(***v***) returns the largest *k* values in ***v***

BatchRefiner(*E,* mode=‘filter’) = *E* × *W_filter_*

**Fig. 2.**
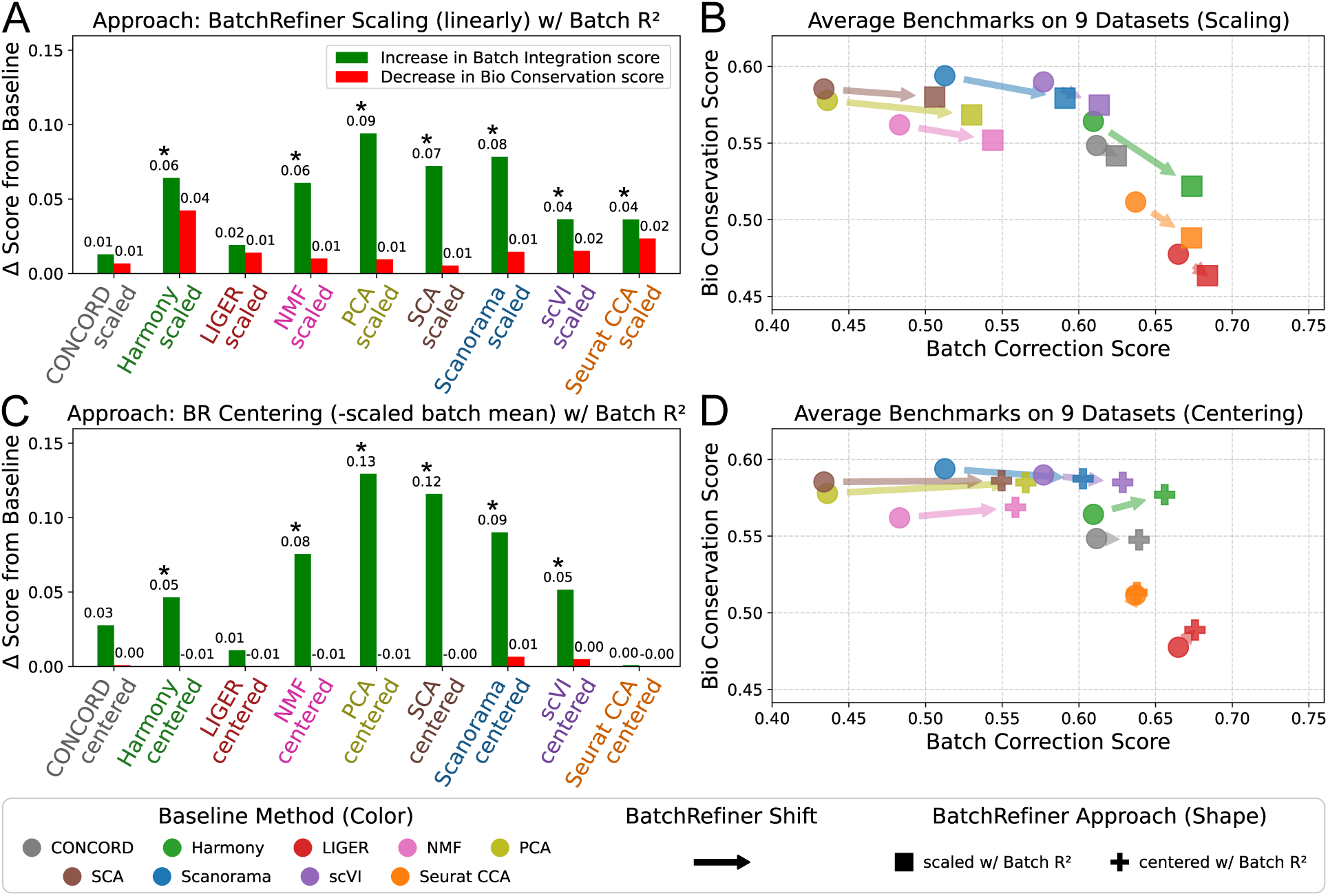
**(A)** Benchmarking BatchRefiner scaling: increases in average batch-integration score (green; * Bonferroni *p <* 0.05) and decreases in average bio-conservation score (red) relative to the corresponding baseline (x-axis) method (100 dimensions, linearly scaled by batch variance explained, or *R*^2^). **(B)** Benchmark values for those nine baseline and BatchRefiner-scaled embeddings, shown as mean bio-conservation (y-axis) and batch-integration (x-axis, Batch Correction) scores, averaged across nine scRNA-seq datasets. Circles, baseline embeddings colored by method as shown in the top right; squares, scaled embeddings. Shifts from baselines to the corresponding scaled embedding are shown with arrows. **(C-D)** Changes in average benchmark scores, as in (A-B), for BatchRefiner (BR) centering (subtracting the batch mean, weighted by batch *R*^2^, from each dimension; plus symbols).

#### BatchRefiner Centering

For centering (by subtracting scaled batch means), we found it generally preserves or even slightly improves bio conservation, which is consistent with its design objective (Figure 2C-D). Centering provides significant increases in batch correction overall (Fisher’s combined *p* = 2.2 × 10^−25^) and for nine individual methods (Bonferroni *p <* 0.05, Supplementary Table 2). In particular, NMF, Scanorama, SCA, and PCA actually have larger increases in average batch correction (0.09 − 0.14) than with scaling (Figure 2A-B), and show essentially no change in bio-conservation scores. Applying BatchRefiner centering to Harmony and scVI yields smaller increases in batch correction (0.05), but still state-of-the-art tradeoffs considering the minimal loss in bio conservation. CONCORD demonstrates a small improvement in batch integration with centering, but the improvement in CONCORD after BatchRefiner post-processing is not statistically significant after Bonferroni correction. LIGER and Seurat, which both operate on initial batch-specific embeddings, show essentially no change in metrics after centering. Again, centering still provides significant overall improvements in batch correction even when the three non-correcting methods are excluded (Fisher’s combined *p* = 1.4 × 10^−14^).

Across the nine individual datasets, we see much variation in the absolute and relative performance of baseline and BatchRefiner-modified embedding methods (Supplementary Figs. S1-S2). This further demonstrates that dataset choice has substantial impact on embedding benchmarks, highlighting the importance of choosing a wide variety of datasets when evaluating new methods as well as the importance of method selection for sensitive downstream tasks. Scaling offers powerful increases in batch correction with no loss in bio conservation on some datasets, such as HypoMap (Supplementary Fig. S1D), as does centering in conjunction with some baseline methods (Extended Data Fig. S2). On the other hand, the magnitude of improvements are much lower for other combinations of method and datasets, such as scaling Harmony and scVI for the DKD dataset (Supplementary Figure S1B).

#### BatchRefiner Filtering

We next considered filtering, removing the lowest scoring fifth, third, or half of dimensions from each embedding (Supplementary Figs. S3-S6, Supplementary Results). In general, we found that filtering has the potential for large increases in batch correction for several baseline methods, but causes larger declines in bio conservation, which were prohibitive for some methods. As expected, the magnitude of effects increased when filtering out more dimensions. Observing that the number of dimensions filtered has a substantial effect of performance and varies with method and dataset, we next evaluated three approaches for filtering a variable number of dimensions based on a fixed score threshold or distribution property (Methods, Supplementary Table 3). However, we did not find that any variable-filtering approach produced consistently better benchmark tradeoffs (Supplementary Fig. S7, Supplementary Results).

We provide a qualitative example of the effect on batch integration and bio conservation of the three BatchRefiner modes in Fig. 3, which shows UMAP plots [45] for Scanorama alone and with BatchRefiner for the Immune Cell Atlas dataset, and for Harmony along and with BatchRefiner for the Tabula Muris dataset. Unlike scaling and centering, filtering displays a clear disruption of some coherent cell types, though also visibly improved batch mixing—strikingly so for Tabula Muris (Harmony). Consistent with the benchmarks (Supplementary Figs S1-S2E,H), scaling and centering on both baseline embeddings show more subtle increases in batch mixing, without visual effect on cell type separation.

**Fig. 3.**
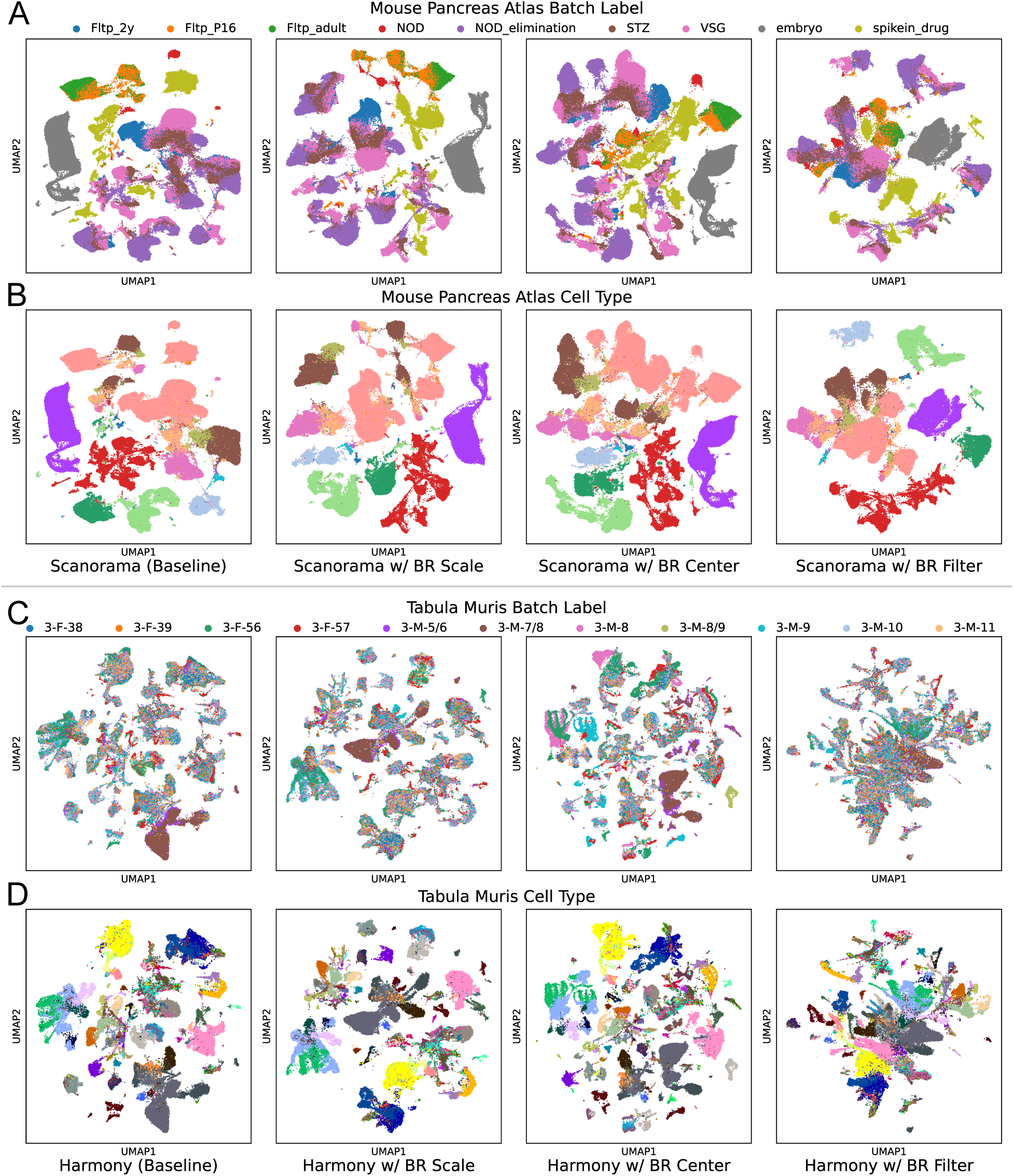
UMAP visualizations of selected scRNA-seqembeddings. **(A-B)** Embeddings of the Immune Cell Atlas dataset colored by **(A)** batch (batch labels shown at top) and **(B)** cell type. From left to right, the UMAP is based on Scanorama alone and with BatchRefiner (BR) scaling, centering, and filtering (one third of dimensions) using batch *R*^2^. **(C-D)** Embeddings of the Tabula Muris dataset colored by **(C)** batch and **(D)** cell type. From left to right, the UMAP is based on Harmony alone and with BatchRefiner scaling, centering, and filtering using batch *R*^2^.

#### Other Variants of BatchRefiner

We also investigated the effect of varying the number of initial embedding dimensions (Supplementary Data). We found that decreasing from 100 to 50 dimensions had a very small negative effect on the batch-correction performance of baseline methods, a minimal effect on BatchRefiner scaling (Supplementary Fig. S8A-B), and reduced the bio-conservation losses for filtering some baseline methods (one third of dimensions, Supplementary Fig. S8C-D). We also found that increasing the initial embedding dimension to 200 for four methods had little effect on baseline and scaled embedding benchmarks, but increased bio-conversation losses from filtering (Supplementary Fig. S8E-F). We also note variation in per-metric effects across metrics (Supplementary Fig. S9) and datasets (Supplementary Figs. S10-SS12, Supplementary Data).

We see similar trends across all nine datasets when using iLISI scores in place of batch *R*^2^ for BatchRefiner, even though the former is a measure of local (cell-neighborhood-based) batch integration and the latter is a measure of global (correlation-based) integration. We see generally similar results from scaling with *R*^2^ (Fig. 2B-C, Supplementary Fig. S1) and with iLISI scores (Supplementary Figs. S13A-B,S14). Using iLISI for centering (Supplementary Figs. S13C-D,S15) actually produced larger increases in batch correction for some methods compared to *R*^2^ (Figs. 2D-E, Supplementary Fig. S2), while still maintaining bio conservation. On the other hand, using iLISI for filtering led to generally larger average losses in bio conservation (Supplementary Fig. S16) than using *R*^2^ (Supplementary Fig. S3), with the notable exception of LIGER. We also observed similar (small) effects of varying the number of initial embedding dimensions when using iLISI scores (Supplementary Fig. S17).

### Benchmarking of scATAC-seq Embeddings

#### scATAC-seq data

While scRNA-seq profiles recently-transcribed genes, it does not reveal any information about genome organization or the regulatory processes upstream of transcription. In contrast, scATAC-seq, which profiles chromatin accessibility, can reveal genomic sites that are bound by proteins that regulate gene expression and chromatin conformation. Generally, DNA that is not actively bound by nucleotide-binding proteins or complexes will be found tightly coiled around histones and inaccessible. Thus, mapping the ATAC-seq sequencing reads back to the genome reveals approximately regions that are actively bound or accessible for binding. Any of several peak-calling approaches can be used to identify a set of genomic regions to which a significant number of reads map; alternatively [20], a pre-defined set of possible enhancers, such as ENCODE canonical *cis*-regulatory elements (cCREs) can be used [46]. Then, for single-cell ATAC-seq, reads are—like scRNA-seq—associated with a particular cell, leading to a cell-by-peak or cell-by-element matrix. Because peak sets typically number in the hundreds of thousands, and because there are only two copies of a chromosome (excluding aneuploidies) versus potentially more transcripts of a gene, scATAC-seq matrices tend to be even sparser than scRNA-seq matrices. We benchmarked a group of widely-used scATAC-seq embedding methods to test their performance in both bio conservation and batch correction.

#### Current scATAC-seq methods

In contrast to methods for scRNA-seq data, there are an assortment of methods used for generating *unintegrated* embeddings—reflecting the greater complexity of embedding scATAC-seq data—but comparatively little variety in the use of batch-correction methods. We selected three widely-used embedding methods to evaluate: pyCistopic [47, 48], Snap-ATAC2 [49], and Signac [50]. None performs intrinsic batch integration, but all methods heuristically suggest—via documentation—the use of Harmony, which was developed for integration of PCA embeddings of scRNA-seq [25]. We therefore benchmark these three with and without Harmony. We separately benchmark Signac in combination with Seurat for batch integration, which is the default option (instead of Harmony) as Signac is integrated into the Seurat ecosystem. We also selected peakVI, a variational autoencoder for scATAC-seq data in the style of scVI that *does* incorporate a batch correction objective [51]. Finally, as a baseline we incorporated the vanilla latent semantic indexing (LSI) procedure [22], an approach adapted from language processing that combines topic modeling with straightforward singular value decomposition, which we refer to as SVD. We also evaluated SVD with Harmony.

We extended the OpenProblems pipeline and metrics to accommodate embeddings of arbitrary single-cell modalities, which allows us to apply the same metrics as used for scRNA-seq to our selected scATAC-seq embedding methods and datasets. There has been little work on benchmarking batch integration specifically of scATAC-seq—one recent work introduced a pair of different metrics to benchmark ATAC-based embedding of multiomic (RNA+ATAC) single-cell datasets, using data from two studies [24]. We assembled a panel of nine public scATAC-seq datasets (Methods), including the largest dataset from each of the two studies previously used for scATAC-seq benchmarking (the Cheong and Weinand datasets, Methods). Each of our datasets is based on a separate *de novo* peak matrix, not a fixed set of cCREs. For convenience, we label all scATAC-seq datasets by first author. We then implemented and provide scripts to filter, adjust cell labels, and unify dataset formatting for benchmarking (Methods).

We observe that the batch-corrected methods clearly improve average integration, with generally no penalty on bio conservation relative to their unintegrated intermediates (Table 2, Figure 4). Overall, pycisTopic and SnapATAC2 both augmented by Harmony appear to offer the best overall performance, with batch integration scores of 0.57 and 0.55, respectively, and bio-conservation scores of 0.61. This is consistent with a previous smaller-scale study using a unique benchmark, which found SnapATAC2+Harmony to be top-performing among a set that did not include pycisTopic or the earlier implementation cisTopic [24]. Among the two datasets shared with previous benchmarking, we find that pycisTopic+Harmony slightly outperforms SnapATAC2+Harmony on the Cheong dataset (Figure 4B), and offers slightly better integration but slightly worse bio conservation on the Weinand dataset (Figure 4I). On our overall benchmarking, SVD+Harmony shows competitive batch integration (0.55) but reduced average bio conservation (0.57), while peakVI has excellent bio conservation (0.61) but a bit less batch integration (0.49; Table 2). Both SVD+Harmony and peakVI can be found among the top methods for some datasets (Figure 4). Finally, we observe that Signac, which is based on LSI, has substantially worse bio conservation than all other initial embedding methods, around 0.47 compared to 0.56-0.59 for others. Both Harmony and Seurat offer improvements when separately run on Signac, with Seurat providing less integration (0.56 versus 0.59 for Harmony) but slightly improving bio conservation (0.50 versus 0.48 for Harmony, Table 2). This is in contrast to our observations on scRNA-seq data for Harmony and Seurat (both PCA-based there), where Harmony better preserved bio integration but did not integrate quite as much as Seurat. The tradeoff we quantified between Signac+Harmony and Signac+Seurat should help users choose in which way to use Signac depending on whether batch integration or bio conservation is more important in the downstream task.

**Table 2.**
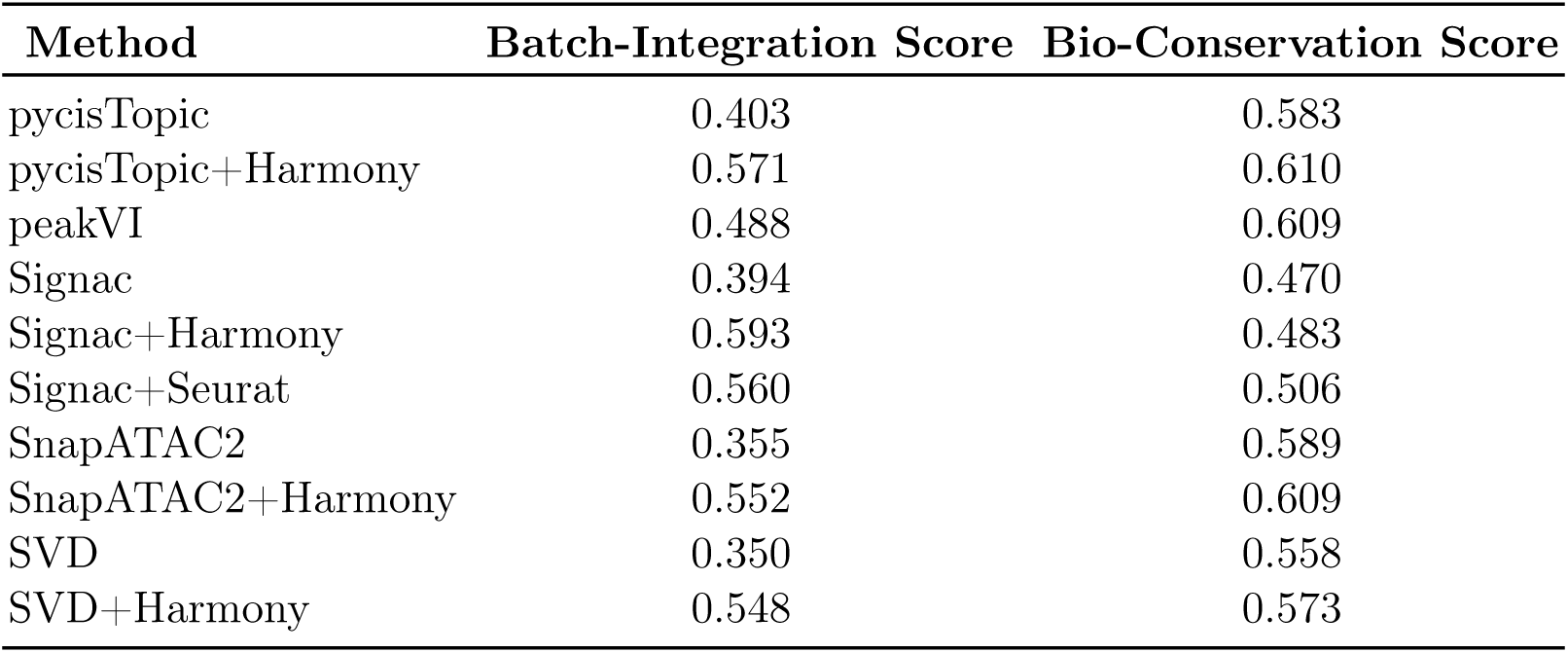
Benchmarking of scATAC-seq embedding methods on nine datasets.

**Fig. 4.**
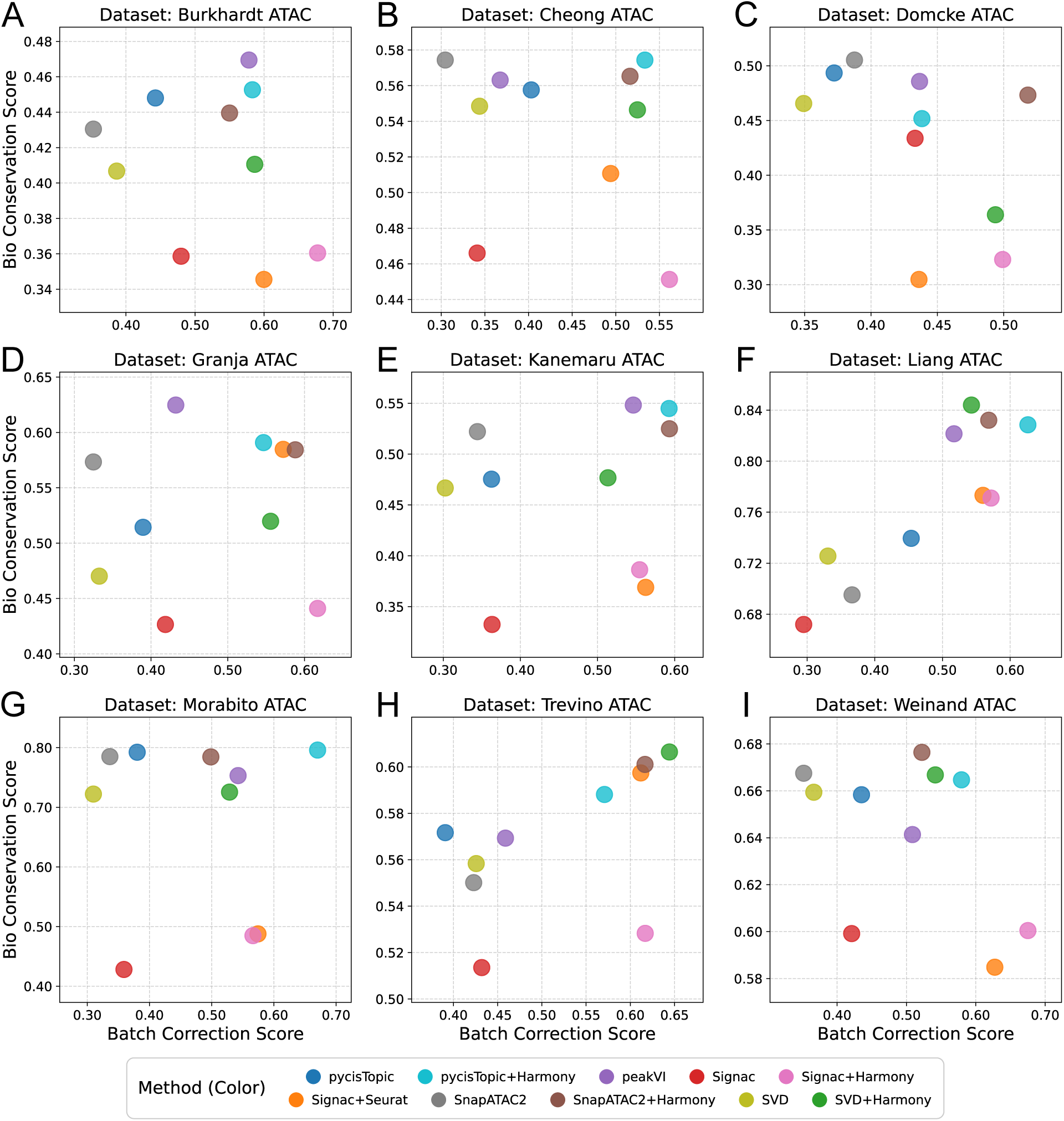
Benchmarks of scATAC-seq embedding methods on each of nine datasets. Benchmarking of embedding methods, shown as bio-conservation (y-axis) and batch-integration (x-axis, Batch Correction) scores for each of nine scATAC-seq datasets individually. These scores are the average of six and five, respectively, individual metrics that are each scaled per-dataset (Methods). Benchmark values range from 0 to 1, with 1 best. Datasets are labeled by first author (see Methods). We did not measure peakVI with Harmony because peakVI incorporates a batch integration objective.

#### Applying BatchRefiner to scATAC-seq

We then applied the three BatchRefiner modes, scaling, centering, and filtering, to these 10 sets of scATAC-seq embeddings. Beginning with scaling using batch *R*^2^ (Figure 5A-B, Supplementary Fig. S18), we observe an overall improvement in batch integration (Fisher’s combined *p* = 2.1 × 10^−24^) and significant increases in batch correction for seven of ten individual baseline methods (Bonferroni *p <* 0.05, Supplementary Table 4). Unsurprisingly, we see large improvement for the four non-integrating methods (pycisTopic, Signac, SnapATAC2, and SVD). The best overall batch integration is achieved by scaling SVD+Harmony, with an increase of 0.08 leading to a score of 0.63; but, this is accompanied by a decrease in bio conservation. Thus, the best combined performance is still offered by pycisTopic+Harmony and SnapATAC2+Harmony, for which BatchRefiner scaling achieves small increases in batch integration (to 0.60 and 0.61, respectively), and accompanied by only minimal reduction in bio conservation to 0.59 (Figure 5A-B). For just the six batch-integrating methods, BatchRefiner scaling still significantly increases batch integration overall (Fisher’s combined *p* = 6.3 × 10^−10^).

**Fig. 5.**
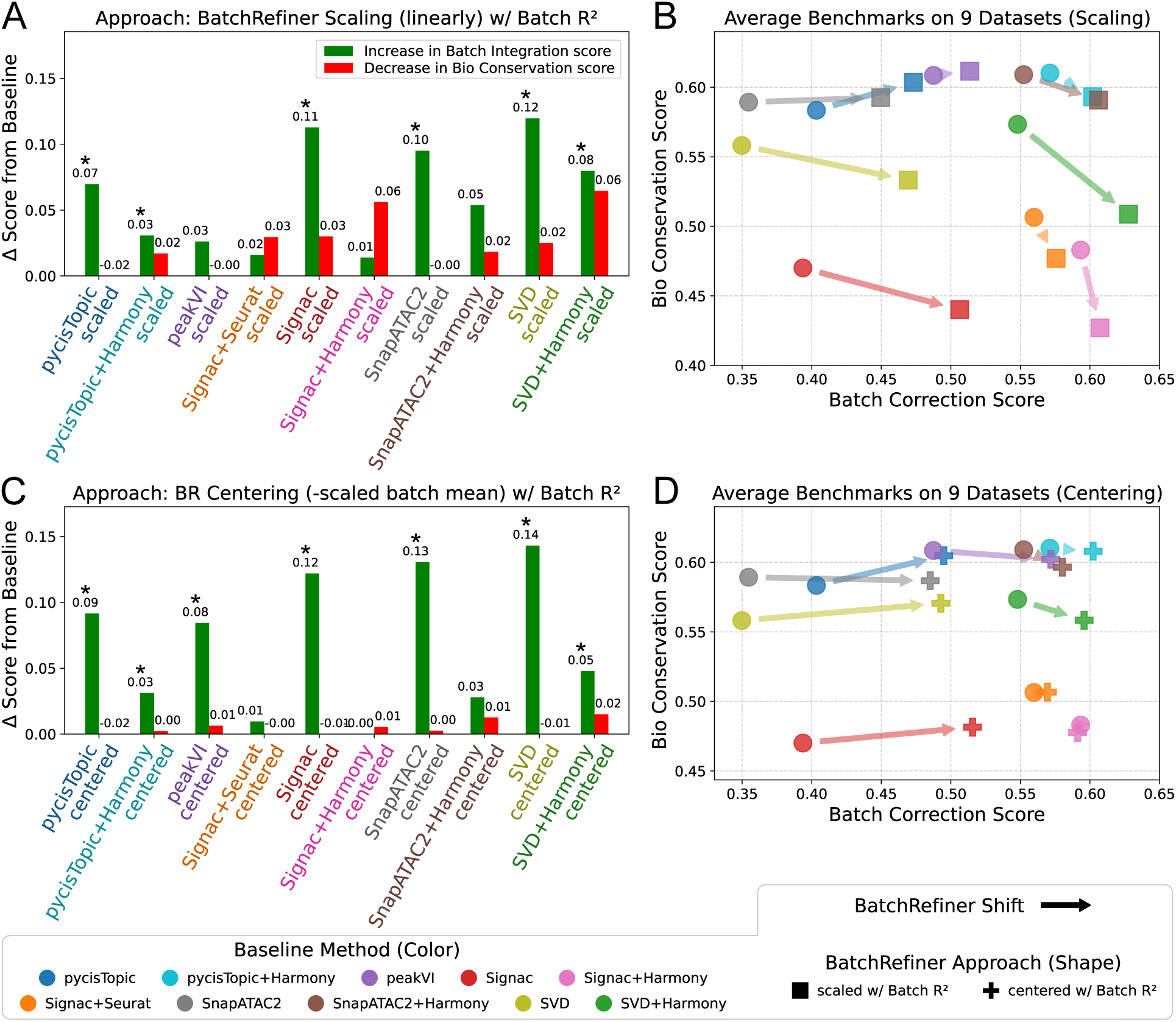
**(A)** Benchmarking BatchRefiner scaling on scATAC-seq: increases in average batch-integration score (green; * Bonferroni *p <* 0.05) and decreases in average bio-conservation score (red) relative to the corresponding baseline (x-axis) method (100 dimensions, linearly scaled by batch variance explained, or *R*^2^). **(B)** Benchmark values for those nine baseline and BatchRefiner-scaled embeddings, shown as mean bio-conservation (y-axis) and batch-integration (x-axis, Batch Correction) scores, averaged across nine scATAC-seq datasets. Circles, baseline embeddings colored by method as shown in the top right; squares, scaled embeddings. Shifts from baselines to the corresponding scaled embedding are shown with arrows. **(C-D)** Changes in average benchmark scores, as in (A-B), for BatchRefiner (BR) centering (subtracting the batch mean, weighted by batch *R*^2^, from each dimension; plus symbols) applied to scATAC-seq embeddings.

For centering (Figure 5C-D, Supplementary Fig. S19), we observed even greater increases in the batch integration of the four non-integrating methods, with no loss or even a slight improvement in bio conservation. And, centering does significantly increase batch integration overall (Fisher’s combined *p* = 2.9 × 10^−29^), when restricted to the six batch-integrating methods (Fisher’s combined *p* = 2.8 × 10^−10^), and for seven of ten individual methods (Bonferroni *p <* 0.05, Supplementary Table 4). Here, the best overall result is achieved by centering pycisTopic+Harmony; this (like scaling) has a small, but significant increase in batch integration to 0.60, but with no loss in bio conservation. On the other hand, the improvements in snapATAC2+Harmony and SVD+Harmony are smaller than those observed from scaling (Figure 5). Centering has no effect on the two batch integrations of Signac (Seurat or Harmony), which is consistent with the observed behavior of scaling on scRNA-seq methods that begin with dataset-specific embeddings (Seurat, LIGER). Finally, peakVI realizes a comparatively large batch-integration increase of 0.08 with centering, with negligible loss of bio conservation, which brings the combined performance in line with the top methods (batch-integration and bio-conservation score of 0.57 and 0.6, respectively).

Lastly, we applied filtering (one third of dimensions) to the scATAC-seq embeddings. Unsurprisingly, we again observed very large loss in bio conservation (Supplementary Figs. S20-S21). Consistent with the observed benchmark of Harmony for scRNA-seq, the four Harmony-based methods here show gains in batch correction of at most 0.06, with loss in bio conservation of 0.14-0.32. One exception is the baseline pycisTopic, which has an increase in batch integration of 0.12 with a loss in bio conservation of only 0.01; but, the combination of pycisTopic+Harmony still shows better performance than pycistopic+filtering alone, and suffers a loss in bio conservation from filtering in addition to Harmony. Our results (loss in bio conservation) from filtering one third of the scATACseq embedding dimensions seem generally more similar to filtering one *half* of scRNA-seq embedding dimensions, which could suggest that scATAC-seq embeddings contain less dispensable information overall.

## Discussion

We presented BatchRefiner, a general, quick post-processing tool based on ensemble refinement, for single-cell embedding methods. Our standalone BatchRefiner Python package supports arbitrary single-cell embedding inputs. The input can be either array data, such as a numpy or scipy object, or the AnnData object [52] used by scanpy [11], which is part of the broader scverse suite of analysis packages [12]. This compatibility enables seamless integration with varied downstream single-cell analyses. The BatchRefiner package also accepts any user-supplied functions for dimension scoring, in addition to built-in support for batch *R*^2^ and iLISI, the two scoring methods we used for developing and evaluating BatchRefiner. The runtime of the BatchRefiner package on a 100-dimensional Scanorama embedding of Tabula Sapiens, the largest scRNA-seq dataset (483k cells), is **12 seconds** when using *R*^2^, or 14 minutes when using iLISI (running on a 64-core server with 2 AMD Epyc 7502 CPUs).

BatchRefiner’s flexibility is such that it could be used on top of any embedding-based batch integration tool, including popular methods such as Scanorama [33, 34], Harmony [25], and scVI [36]. Furthermore, the approaches of BatchRefiner—scaling, centering filtering—yield different tradeoffs between batch integration and bio conservation, allowing users to select the preferable mode for each dataset and downstream application. We show examples of this phenomenon using individual benchmarks, many of which are proxies for common downstream tasks; for example, the cLISI (cell-type LISI) bio-conservation metric is a proxy for *k*-nearest-neighbor cell classification [25]. We also show a visual example of increasing batch integration using UMAP (Figure 3) of scRNA-seq embeddings, a common downstream visualization and itself a proxy for other locality-based tasks.

We presented the entire scRNA-seq benchmarking and analysis first because the benchmarking framework that we generalize [16] was established for scRNA-seq. In summary, the best method by average performance on our nine scRNA-seq datasets was Harmony combined with the BatchRefiner centering approach. However, each embedding method can have useful properties for specific datasets and tasks beyond simply what is captured by the bio-conservation and batch-integration benchmarks. This observation motivated our robust benchmarking of many methods and our consideration of BatchRefiner applied, with multiple approaches, to all these methods. When our analysis is extended to scATAC-seq methods and nine scATAC-seq datasets from separate studies, the overall best method is pycisTopic with Harmony and BatchRefiner centering; pyCistopic or Snap-ATAC2 with Harmony and BatchRefiner scaling also have very strong average performance. The benchmarking framework extension to scATAC-seq with current methods yields cutting-edge benchmarking results, independent of the addition of BatchRefiner; **therefore, our pipeline can be used to robustly and independently evaluate newly-developed scATAC-seq embedding methods.**

We observe that the tradeoff between batch integration and bio conservation applies across multiple datasets and tools. As in previous benchmarking efforts [8], we find that the best scRNA-seq methods usually appear on a “tradeoff frontier”, represented by a negatively-sloped line, similar to a Pareto frontier. We do not observe such a frontier for scATAC-seq, as the top methods show similar batch-integration performance, which may be attributable to the majority of batch-correcting approaches we benchmarked being Harmony-based. Nonetheless, BatchRefiner achieves state-of-the-art performance in both the scRNA-seq and scATAC-seq modalities by providing larger gains in batch integration than what would be expected by simply moving along the frontier to a state with a lower bio-conservation score.

For the two BatchRefiner approaches that we highlight, scaling and centering, we have focused only on the default option of a linear scaling function. This demonstrates the potential of BatchRefiner’s ensemble refinement approach without parameter tuning, and thus without overfitting. And, when reporting average benchmarks, all method parameters were shared across all datasets. While the group of top-performing methods is generally shared across datasets, the observed performance of individual embedding methods can vary substantially between datasets. Future work could investigate the role of ensemble refinement in designing dataset-specific embedding workflows [18].

As we demonstrated, our stacked generalization approach to modify embeddings (Box 1) suffices to statistically, albeit empirically, improve batch integration across a wider range of methods as applied to a recent, comprehensive set of benchmarking datasets and benchmarks established by others in the field [16]. Further, our scaling and especially centering approaches improve batch integration with little to no loss in bio conservation, which measures–among other properties— the preservation of distinct cell types in the integrated embedding. That said, they do not offer a guarantee that they generate the optimal reweighting or filtering of an embedding space. Moreover, BatchRefiner considers each embedding dimension independently, which could obscure batch effects observable only jointly in multiple dimensions, especially for embedding methods that may produce correlated dimensions. On the other hand, this simplicity is key to BatchRefiner’s flexibility and speed. Because BatchRefiner makes no assumption about the embedding structure, it could be applied to batch-integrated embeddings of single-cell modalities other than scRNA-seq and scATAC-seq.

## Conclusions

The BatchRefiner tool for ensemble refinement significantly improves scRNA-seq and scATAC-seq batch integration across several state-of-the-art embedding methods applied to a comprehensive set of datasets and benchmarks established by others in the field [16] for scRNA-seq, addressing an ongoing challenge in robust analysis of atlas-scale data [17]. Because BatchRefiner takes a cell embedding matrix as input, it is method-independent and generalizable to any current or future cell embedding method. Furthermore, our strategy to assess significance can be applied to any pair of methods for any single-cell task, and a Fisher’s combined test, to any post-processing method run on a set of baseline methods. Multimodal single-cell assay technologies contribute orthogonal biological information [30–32], and multiomic data continue to grow [5]. As these reach atlas scale and related embedding methods are developed, ensemble refinement is innately generalizable to additional omics modalities. We expect BatchRefiner’s fast, plug-and-play approach for ensemble refinement of embeddings, along with its significance statistic for improvement in batch integration, to be integral parts of many single-cell analysis toolkits.

## Methods

### scRNA-seq Datasets for the OpenProblems Benchmarking Pipeline

We utilized the Batch Integration task (version 2) from the OpenProblems framework [16] for benchmarking embeddings of scRNA-seq datasets. The framework includes two pre-processing steps. The first step processes input data into a “common” (OpenProblems-wide) format, including standardizing the data naming and performing several normalization methods. The second, task-specific pre-processing step performs additional operations, in particular restricting genes to 2,000 batch-aware highly variable genes (HVGs) in each dataset via scanpy [11]. The OpenProblems website uses six datasets from CELLxGENE [5] for this pipeline; we downloaded the fully processed datasets from [53]. The datasets included are: DKD (human Diabetic Kidney Disease, 39,176 cells) [54]; GTEx v9 (human cross-tissue, 209,126 cells) [55]; HypoMap (mouse hypothalamus, 384,925 cells) [56]; Immune Cell Atlas (human, 329,762 cells) [57]; Mouse Pancreas Atlas (pancreatic islets, 301,796 cells) [58]; and Tabula Sapiens (human cross-tissue, 483,152 cells) [6]. So that our results on these datasets would cleanly extend OpenProblems, we did not modify these datasets at all.

We augmented these datasets with three additional scRNA-seq datasets to increase the power and variety of our benchmarking. To accommodate these, we slightly extended the OpenProblems data processing pipeline to support adding arbitrary new datasets, outside of CELLxGENE, in AnnData format. We obtained an embryonic developmental atlas of *C. elegans* cells [59] from NCBI GEO accession GSE126954; this dataset was previously used in recent batch-integration benchmarking [40]. For this dataset, we used the provided batch labels. But, we redefined the cell type annotations based on the original authors’ supplementary comments to reconcile two incomplete groups of labels (‘cell.type’ and ‘cell.subtype’) into a consistent set. We kept 62,178 cells to which we could assign clear terminal cell-type labels, which excludes early embryonic cells, totally unannotated cells, and cells marked as failing quality control.

The next dataset is the Tabula Muris dataset, a pan-tissue mouse atlas [60], which we obtained from Figshare (datasets 5968960 and 5829687 for two different scRNA-seq technologies, Droplet and FACS). This dataset was also used in a previous benchmarking study [15]. Here, we defined the batch labels as the cartesian product of the mouse identifier and the experimental technology used (Droplet or FACS). We took the provided cell labels in the ‘cell_ontology_class’ field, excluding unlabeled cells, except that we also included cells free-labeled with ‘chondrocyte-like’. We also filtered out two batches with fewer than 100 cells each. This resulted in a final total of 99,915 cells. Finally, we obtained an atlas of the human eye (Ocular Atlas, 128,016 cells) from the Single Cell Portal accession SCP2310 [61]. Here, we simply used the patient identifier as the batch and took the provided coarse cell type labels in the ‘class’ field.

### Benchmarking scRNA-seq Embedding Methods via OpenProblems

Using the Batch Integration task (v2, as of October 2025) pipeline, we benchmarked baseline and BatchRefiner methods on those nine datasets. We modified the benchmarking pipeline as required to run on a local machine via docker [62]. We added or modified each baseline method into the benchmarking pipeline. Harmony (using the harmonypy implementation version 0.0.10 [63]), scVI (using scvi-tools [37] version 1.4.0), and LIGER (using the rliger package [39]) were already implemented in the OpenProblems framework; we modified these to produce 100-dimension outputs for consistency, instead of the default 50, 30, and 20 dimensions, respectively. LIGER has a maximum dimension equal to the smallest batch size, which was between 20 and 100 for two of six datasets (see below). While doing our initial benchmarking with the OpenProblems framework, we recognized that OpenProblems erroneously evaluates Scanorama using an undocumented combination of Scanorama’s embedding and count-correction outputs, which are separate outputs of inherently different types. Therefore, we modified the pipeline to benchmark the Scanorama embedding output only. Our correction to the Scanorama usage, as well as our addition of SCA, have subsequently been merged into the OpenProblems batch integration repository.

We added the remaining five baseline methods. We used the Scanpy [11] implementation of PCA and the scikit-learn [43] implementation of NMF. We ran SCA using the original package [42], running with 5 iterations as suggested. We used the Seuratv5 implementation [23] of canonical correlation analysis (CCA, following [64]) version 5.3.0. Finally, we also ran CONCORD using the original package [40], and all genes. We initially specified 100-dimension outputs and otherwise used default parameters. For Seurat CCA, the k.weight was set to the minimum of 100, the default value, and the number of cells in the smallest batch, which is the maximum allowed value. Finally, we adapted scVI, Seurat, and CONCORD to have the option to load in embeddings computed (using the same implementation) outside of the pipeline. In particular, this addressed GPU (scVI, CONCORD) and time (Seurat) requirements that were difficult to accommodate in our local pipeline deployment. When benchmarking precomputed embeddings, we manually placed intermediate outputs in the pipeline’s working directories. We note that scVI and CONCORD train machine learning models (a variational autoencoder and a “minimalist” neural network, respectively) to embed each dataset, and this approach ensured that the same scVI and CONCORD models/embeddings were used across all baseline and BatchRefiner runs for each dataset and embedding dimension. After benchmarking 100-dimension embeddings from each method, we subsequently benchmarked 50-dimension embeddings from all nine methods and 200-dimension embeddings from CONCORD, Harmony, Scanorama, PCA, and SCA.

### Benchmarking Metric Calculations

The OpenProblems batch-integration benchmarking task uses 12 metrics, 5 for batch integration and 7 for bio conservation, to compute per-methods scores. The batch-integration metrics are i. principal component regression (PCR) comparison, ii. batch-mixing average silhouette width (ASW), iii. graph connectivity score, iv. graph iLISI, and v. k-nearest neighbor batch effect test (KBET). The bio-conservation metrics are i. isolated label ASW, ii. k-means normalized mutual information (NMI), iii. k-means adjusted Rand index (ARI), iv. cell-type mixing ASW, v. cell-type LISI (cLISI), vi. cell cycle conservation score, and vii. isolated label F1 [8, 16]. We highlight that several of these metrics evaluate how cells that share the same cell type label group together, both locally (*e.g.*, cLISI) and globally (*e.g.*, cell-type ASW). OpenProblems’ results also include a 13th metric, highly-variable gene overlap, that is not run for embedding methods, which we therefore skipped when computing mean scores. We also excluded from the per-dataset averages any missing metrics; in particular some datasets lack any “isolated” cell-type labels(Supplementary Figs. S10-S12F,K). (This diverges from the behavior of the OpenProblems website, which treats missing metrics as 0 values.) Each metric modifies the underlying score, if needed, to have a range of 0 to 1, with 1 being theoretically perfect, although not necessarily achievable for a real dataset [16].

In practice, different metrics have different ranges of achievable scores. We therefore followed the strategy of OpenProblems in computing reference ranges using a panel of seven “control” methods, including methods that produce random embeddings and that directly embed cell type information. Because some include randomness, we benchmarked each control method six times (instead of only once as in OpenProblems) and took the union of metric values. We also corrected the behavior of the “embed_cell_types” control method, to match its definition, by removing the inclusion of random noise to the cell type embeddings used. (There is also corresponding control method *with* random noise.) We then re-scaled the scores of each metric in each dataset by mapping the minimum and maximum values achieved by any control method to 0 and 1, respectively. Metric values for non-control methods that fall outside of the reference range are reassigned to be 0 or 1. Thus, when subsequently averaging multiple metrics to produce aggregate batch-correction and bio-conservation scores, the values of each metric are normalized to represent the achievable range of values, and therefore each metric is expected to contribute approximately equally to the multi-metric average. We note that the panel of control methods typically provide a wider range of performances than any plausible method would. So, empirically the reference range is still cautiously large and several individual methods tend to produce re-scaled values that fall in smaller windows (Supplementary Figures S10-S12, Supplementary Data). However, the alternative of re-scaling based on observed performance of “real” methods would result in a re-scaling specific to each benchmarking that would not be comparable across method sets. Further, this alternative would be overly restrictive given that all methods perform some degree of interpretation and bio-conservation and that it remains possible that newly-developed methods, such as BatchRefiner, will substantially outperform the current range of achieved values for some metrics.

We left all metric implementations in the OpenProblems pipeline unmodified with one exception. We observed very high memory and time usage for KBET (one of several pipeline metrics implemented in the scib package v1.1.7 [8]), in excess of 2TB and 24 hours for certain embeddings of the HypoMap dataset. We note that the public OpenProblems benchmarking results omit (*i.e.*, assign a score of 0 to) KBET for embeddings of HypoMap. We modified the data structure usage in a deterministic preprocessing component to reduce resource usage to 2 hrs and 600GB in the worst observed case, and to generally improve runtime and memory usage. Accordingly, we ran KBET on some candidate embeddings outside of the pipeline, and later integrated our modification into the pipeline. Our improved KBET is available in our fork of the scib package [65] and was subsequently merged into sciv v1.2.0.

We did not add any new metrics to the pipeline, nor did we change which metrics are used to compute average scores. In particular, we computed (Supplementary Data), but did not include in our averages, values of a few additional metrics that appear to have been added to the pipeline subsequent to the generation of the published results.

### Benchmarking of BatchRefiner modes on scRNA-seq data

For each of the nine embedding methods, we created several BatchRefiner-postprocessed copies of each method to scale, center (subtract scaled means), and filter (parameterized as to the number of dimensions to filter), using batch *R*^2^ or iLISI as dimension scoring metrics (below). For PCA, SCA, Scanorama, and Harmony, each copy ran the method *ab initio*, with a parameterized number of dimensions, and scored and post-processed these dimensions. For Seurat, scVI, LIGER, NMF, and CONCORD, our BatchRefiner versions instead loaded their embeddings, previously computed as an intermediate output of benchmarking those methods at baseline, prior to scoring dimensions. Our Seurat, scVI, LIGER, NMF, and CONCORD implementations for centering and filtering with iLISI additionally loaded in per-dimension iLISI scores computed for scaling. Because the maximum output dimension of LIGER is equal to the number of cells in the smallest batch, it output only 83 dimensions for the Tabula Sapiens dataset and only 40 dimensions for the GTEx dataset. In the former case, we kept 42, 55, and 66 dimensions when filtering out 1*/*2, 1*/*3, and1*/*5 of 100 initial dimensions. In the latter case, we kept 20, 27, and 32 dimensions when filtering out 1*/*2, 1*/*3, and 1*/*5 of 50 or 100 initial dimensions. In addition to implementing filtering a specified number of columns, we also implemented additional filtering versions that filtered dimensions based on a specified batch *R*^2^ threshold and that filtered dimensions with batch *R*^2^ scores greater than the third-quartile value plus a multiple of the inter-quartile range (IQR).

We computed the batch *R*^2^ of each embedding dimension by regressing the values of each dimension against one-hot encoding of each batch label. The output is then the fraction of variance of each dimension (*R*^2^) explained by the one-hot batch labels. We note that this is closely related to the PCR comparison metric [13]. In this metric, both unintegrated RNAseq data and a candidate embedding (or corrected gene expression data) are reduced by PCA to 50 dimensions, and the batch *R*^2^ is calculated for each principal component as described (hence, Principal Component Regression, or PCR). These values are then weighted by the fraction of variance explained by each PC, and summed. Finally, in the Comparison step, the weighted PCR values for the unintegrated data and the embedding are compared to quantify the reduction in batch signal in the integrated embedding (and the result is negated, so a larger score is preferred). For a single dimension, the steps of performing PCA and weighting by variance are meaningless. And, the Comparison step is irrelevant to ranking different embeddings of the same dataset, which includes different dimensions of the same embedding. It follows that for single-dimension “weak” embeddings, our batch *R*^2^ is equivalent to using the PCR Comparison metric. The Comparison step would add a fixed transformation based on unintegrated batch *R*^2^, but this is also moot if the set of metric values are subsequently normalized (scaling, centering) or ranked (filtering a fixed number of dimensions).

For scoring dimensions, we used the scib (v1.1.7) implementations for PCR Comparison and iLISI, which are the same implementations used by the OpenProblems pipeline for evaluating embeddings [8, 16]. Given the simplifications of single-dimension batch *R*^2^ compared to full PCR Comparison, described above, we directly invoked an intermediate function that performs regression of each dimension against one-hot encodings of each batch label using scikit-learn [43]. As input to iLISI, we used Scanpy [11] to compute a neighbor graph from single embedding dimensions, with default settings and following the implementation used by OpenProblems in computing the iLISI metric [16]. We omitted the last step of computing the iLISI metric, which is to normalize by the number of batch labels, as it is unneeded when comparing on a single dataset.

### scATAC-seq benchmarking datasets

We assembled a benchmarking panel of nine human scATAC-seq datasets with publicly available peak matrices and cell type annotations. For all datasets, we used count matrices with *de novo* peak calls from the original studies, as well as provided cell-type annotations. In doing so, we aimed to capture a variety of methodological approaches at the experimental and computational stages. We obtained the Burhkardt dataset of 69,249 bone marrow mononuclear cells from NCBI GEO accession GSE194122; this dataset was initially assembled for a benchmark of tool of multimodal integration [66]. We used the provided batch and cell type labels. We obtained the Cheong dataset of 197,360 peripheral blood mononuclear cells (PBMCs) [67] from NCBI GEO accession GSE196987; this dataset was one of two studies used in a recent benchmarking [24]. We used the provided batch and cell type labels for all cells from both healthy and post-infection patients. We obtained the Domcke dataset of pan-tissue fetal cells [68] from NCBI GEO accession GSE149683, including 15 tissue-specific sub-datasets. We used the donor ID as the batch label and the provided cell type labels, except that we filtered out cells of unknown type, merged a few synonymous labels, and merged labels with and without question marks. There were 697,719 cells after filtering; we then reduced this by a factor of four (to 174,429) using GeoSketch [69].

We obtained the Granja dataset of hematopoetic cells [70] from [71]. We took only cells from healthy patients, and used the provided ‘Group’ annotation as the batch label. We extracted the cluster IDs from the ‘Biological Classification’ field and mapped these to cell types using the information provided in the study’s Supplementary Table 1, excluding clusters with the label ‘Unknown’ label. After filtering, we had 33,819 cells. We obtained the Kanemaru dataset of 139,835 cardiac cells [72] by downloading the ATAC peak matrix from [73]. We used the provided batch and cell type labels. We obtained the Liang dataset of 154,775 retinal cells [74] from [75]. We used the Donor UUID as the batch label and the provided cell type labels. We determined that the provided matrix had been normalized by using 10,000 reads per cell (*i.e.*, dividing each cell’s counts by the number of reads divided by 10,000), followed by log transformation [ln(*x* + 1)]. We further determined that the number of reads used for this normalization corresponded to the number of RNA-seq reads from paired transcriptomic providing (the ‘nCount_RNA’ field) rather than the number of ATAC-seq reads per cell. We reversed this normalization to recover the raw peak matrix.

We obtained the Morabito dataset of 130,418 brain cells [76] from NCBI GEO accession GSE174367. We used the sample IDs as batch labels and the provided cell type labels, and we used cells from both healthy donors and patients with Alzheimer’s Disease. We obtained the Trevino dataset of 31,304 fetal cortex cells [77] from NCBI GEO accession GSE162170. We used the sample ID as batch labels, and mapped the provided cluster annotations to cell types using Supplementary Table S1E of the original study. We used only cells profiled by scATAC-seq only, and did not use the smaller dataset of multiomically-profiled cells. Finally, we obtained the Weinand dataset of 31,547 synovial-tissue cells [78] from “dataset 1” of Zenodo record 15073137 [79]. We note that we used the data as provided by the related benchmarking study [24]. We used the provided sample and phenotype annotations and the batch and cell types labels, respectively.

### Benchmarking scATAC-seq Methods via OpenProblems

We generated baseline, 100-dimensional embeddings on each peak matrix for five scATAC-seq embedding methods. We first converted each dataset to a MatrixMarket-like format for use with both R- and Python-based embedding methods. We then compiled each dataset’s metadata, along with precomputed embeddings (below), into AnnData format for use with the OpenProblems pipeline. We used peakVI from scvi-tools [37] version 1.3.3, the pyCistopic implementation of the cisTopic method [48], snapATAC2 version 2.9.0 [49], and Signac version 1.17.1 [50], all using default parameters to generate 100-dimension embeddings. We additionally pre-computed Seurat batch integrations of the Signac embeddings using rLSI from Seurat version 5.5.1 [23]. As for scRNA-seq, the k.weight was set to the minimum of 100, the default value, and the number of cells in the smallest batch, which is the maximum allowed value. Finally, we computed our baseline latent semantic indexing (LSI) using a vanilla TF-IDF implementation followed by TruncatedSVD from scikit-learn [43], keeping the second through 101st components and recording the variance explained.

We further adapted both pre-processing stages of the OpenProblems pipeline (above) to accept datasets with pre-existing embeddings and metadata but no gene expression data. We then proceeded to benchmark baseline and further-integrated embeddings using the pipeline. We ran Harmony (using the harmonypy implementation version 2.0.0 [63]) as well as BatchRefiner scaling, centering, and filtering (as described above) *ab initio* for each pipeline run. We did not compute the cell cycle conservation metric, which requires expression data for genes that mark cell cycle progression, leaving six bio-conservation metrics. And, we modified the PCR Comparison metric to, instead of PCA, use the first 50 dimensions of our SVD (LSI) embedding, and the corresponding variances explained, as the baseline; this was consistent with our consideration of LSI as a ‘default’ reduction method for scATAC-seq. We similarly applied the seven control methods to the metadata and count matrices of the scATAC-seq datasets. For control methods that produce modifications (shuffles) of the count matrix, we again substituted 50-dimension TF-IDF+SVD in place of the 50-dimension PCA used by default by OpenProblems.

### Assessing statistical significance of BatchRefiner improvements in batch correction

We sought to assess whether BatchRefiner can *significantly* improve batch integration relative to each baseline embedding method. So, for each baseline method and for two selected modes (scaling and centering with batch *R*^2^), we performed paired, one-tailed T-tests between the batch-integration scores (average of five scaled metrics) without and with BatchRefiner on the nine datasets. This gave *p*−values for rejecting each null hypothesis that applying BatchRefiner scaling or centering does not increase the batch-integration score relative to a baseline embedding method. We then performed a Bonferroni correction, multiplying each *p*-value by *n* = 18 for scRNA-seq (nine methods × two modes, Supplementary Table 2) or by by *n* = 20 for scATAC-seq (ten methods × two modes,

Supplementary Table 4). Finally, considering that with *n* = 9 datasets we may be underpowered to identify significant improvements, we applied Fisher’s combined probability test [80] to each group of nine (scRNA-seq) or ten (scATAC-seq) *p*-values to evaluate the overall improvement of each BatchRefiner mode. We also performed a Fisher’s combined test restricted to the six baseline scRNA-seq or scATAC-seq methods that perform batch correction. We corrected the Fisher’s method *p*-values by multiplying each by *n* = 2 modes (scaling and centering) evaluated for significance.

## Declarations

### Ethics approval and consent to participate

All human scRNA-seq and scATAC-seq data used in this study were obtained from public repositories (Methods).

### Consent for publication

Not applicable.

### Availability of data and materials

BatchRefiner is available as a standalone, installable package at https://github.com/schafferde/BatchRefiner. Our version of the OpenProblems batch-integration pipeline with all additional and modified methods is available on GitHub in our fork of OpenProblems’ batch-integration repository: https://github.com/schafferde/task_batch_integration/tree/batchrefiner_reproducibility.

This branch also contains other scripts for preprocessing datasets and for generating figures. Complete benchmarking scores are also available in this repository, and are also made available as supplementary data. Our code for processing the three additional scRNA-seq benchmarking datasets and nine scATAC-seq benchmarking datasets (Methods) is available on GitHub in our fork of Open-Problems’ datasets repository: https://github.com/schafferde/openproblems_datasets/tree/more_scrna_datasets. Individual benchmarking datasets were obtained from publicly-available repositories as described in Methods.

### Authors’ contributions

All authors contributed to the conception and design of the methodology. D.E.S. and D.E. collected preliminary data. D.E.S. and H.K. identified datasets and generated embeddings. D.E.S. primarily performed benchmarking and prepared figures. D.E.S, H.K., E.D.A. and B.B. analyzed and interpreted data. D.E.S and B.B. drafted the manuscript text. All authors revised and reviewed the manuscript. B.B. guided the project.

## Acknowledgments

We thank Maxwell Sherman for helpful discussions.

