## Supplementary Materials for "BatchRefiner: fast, significant improvement in batch integration of single-cell embeddings with ensemble refinement"

### Supplementary Results

#### BatchRefiner Fixed-Dimension Filtering for scRNA-seq

We benchmarked the BatchRefiner filtering mode, filtering out the worst-scoring fifth, third, and half of dimensions using batch $R^2$. This corresponds to removing 20, 33, and 50 dimensions, respectively, from a 100-dimension initial embedding of each scRNA-seq dataset. In general, we observe that filtering can offer larger increases in batch correction; however, the tradeoffs in loss of bio conservation are mixed (Extended Data Fig. 3). As one might expect, the magnitudes of gains and losses generally grow with the number of dimensions removed, but often at different rates from each other.

We take the average scores using the intermediate filtering approach tested (filtering out 1/3 of dimensions, Extended Data Figs. 3C-D,4) as representative. For NMF, SCA, and Scanorama, we observe large increases in batch integration ($0.12 - 0.14$) with smaller losses in bio conservation ($0.05 - 0.07$). In general, for these methods both batch-correction gains and bio-conservation losses are smaller when filtering fewer dimensions (1/5, Extended Data Fig. 3A-B, Supplementary Data Fig. S5) and are larger when filtering more dimension (1/2, Extended Data Fig. 3E-F, Supplementary Data Fig. S6). For PCA, we observe a large average increase in batch integration (0.19) but also a large decrease in bio conservation (0.10); the bio-conservation loss improves to only 0.04 when filtering 1/5 of dimensions (with 0.14 batch integration increase; Extended Data Fig. 3A) and worsens to 0.21 when filtering 1/2 of dimensions (with 0.23 batch integration increase; Extended Data Fig. 3E). Filtering is not effective for LIGER which has high batch correction but low bio conservation at baseline, since it heavily decreases bio conservation (0.16) with only minimal gain in batch correction (0.3). Even when filtering fewer dimensions, the bio-conservation loss is still 0.09 with only 0.03 gain in batch correction. For Harmony, we also observe a much larger decrease in bio conservation (0.24) than the increase in batch integration (0.12), although the latter is substantial viewed in isolation. In general, BatchRefiner filtering as applied to Harmony is likely not a valuable tradeoff given the very high loss in bio conservation, even when considering that the tradeoff improves a bit to $(+0.11/-0.18)$ when filtering fewer dimensions (Extended Data Figure 2). We observe little effect from filtering CONCORD, with an increase of only 0.02 in batch correction, but a corresponding bio-conservation loss of only 0.02. Finally, filtering scVI or Seurat shows smaller increases in batch correction and decreases in bio conservation, generally moving along the frontier in the batch-integration direction (Extended Data Figure 3).

### BatchRefiner Variable-Dimension Filtering for scRNA-seq

We observed that the level of filtering that provided the best apparent tradeoff varied between methods, as discussed above, as well as between scRNA-seq datasets. We further observed that the distributions of batch $R^2$ scores are generally left skewed; that is, most dimensions have low scores with a relatively small tail having high batch-signal (Supplementary Data). We therefore investigated filtering methods that would filter a varying number of dimensions depending on the apparent number of high-batch columns, while maintaining an overall straightforward approach. We first tried filtering all columns with a batch $R^2$ of at least 0.01. For two methods, Harmony and Seurat, this filtered between 1 and 48 columns (Supplementary Table 3). The benchmarks of the resulting embeddings were similar to filtering one-fifth of the dimensions for these methods, and importantly the average loss in bio conservation was not mitigated (Extended Data Figure 5A-B). Further, for several other methods, this threshold proved to be too low as most or all dimensions were filtered for most datasets (Supplementary Table 3). (The main exception was HypoMap, which was the same dataset for which only 1 dimension was filtered for Harmony and Seurat.) For evaluation, we had capped the number of dimensions filtered at one half, and unsurprisingly the resulting average benchmarks are very similar to those for filtering one-half of columns (Extended Data Figs. 3E-F,5A-B).

Given this result, we next raised the batch $R^2$ threshold to 0.10. Now, more than 50 dimensions were filtered out in only two cases (SCA and NMF for Mouse Pancreas Atlas; Supplementary Table 3). On the other hand, no dimensions were filtered for HypoMap with any method, for Seurat for seven of nine datasets, or for Harmony for five of nine datasets (Supplementary Table 3). So, for those methods the change in benchmarks, like the change in embeddings, was very small (Extended Data Figure 5C-D). For some methods, namely NMF, PCA, SCA, and Scanorama, there was some increase in batch correction (0.08-0.11) with lower loss in bio conservation (0.02-0.06), but these average results are comparable to those obtained by filtering out a fixed fraction of 1/5 of dimensions (Extended Data Figure 2A-B,5C-D). Given these findings, we concluded that the score distribution and magnitudes vary with both dataset and method, and therefor setting a fixed score threshold is not generally superior to setting a fixed number of dimensions for filtering.

Finally, we investigated a form of outlier detection. We applied a test for (weak) outliers, the third quartile plus the inter-quartile range, and filtered dimensions with batch $R^2$ exceeding that threshold. (The canonical standard for outlier for detection would be 1.5 times the IQR, which is why we refer to this as testing for weak outliers.) This approach filtered between 1 and 17 columns, with a median of 10 (Supplementary Table 3); by definition, it must filter less than 25. In general, the average results again appear similar to filtering 1/5 of dimensions, with slightly lower effect sizes consistent with filtering fewer dimensions (Extended Data Figure 3A-B,5E-F). In particular, Harmony still has a high loss in bio conservation (0.16) to go along with its increase in batch correction (0.10). There are a few methods for which the loss in bio conservation is reduced to negligible - SCA, PCA, and scVI with this IQR-based filtering have only 0.01 loss in bio conservation (compared to 0.02, 0.04, 0.02) while retaining batch-correction improvements of 0.07, 0.12, and 0.03. However, these are still not improved over what is offered by centering for these methods (0.02, 0.13, 0.05; no loss in bio conservation). As the overall benchmarking results are still mixed, we cannot conclude that IQR/outlier-based filtering is generally better at filtering the "correct" number of dimensions. Nonetheless, it may be possible to develop more complex heuristics for filtering that take into account the distribution of scores, and additionally a manual inspection of the score distribution may prove useful for setting a fixed quantity or threshold for a new dataset. To fairly evaluate filtering without overfitting, we did not manually choose dataset-specific thresholds, nor did we consider the specific score distributions when initially choosing 1/5, 1/3, and 1/2 as the fractions of dimensions to filter.

### Supplementary Data

The Supplementary Data file contains thirteen sheets. The first four sheets contain unscaled metric values output by the OpenProblems pipeline for all benchmarking conducted (Supplementary Table S1); there is one sheet each for 50-, 100-, and 200-dimensional initial scRNA-seq embeddings, and one sheet for 100-dimensional scATAC-seq embeddings. The fifth and sixth sheets contains metric values output for 42 benchmarks of control methods (7 methods, 6 replicates) across the nine scRNA-seq datasets and scATAC-seq datasets, respectivly. Next, scaled metric values (see Methods) are in the seventh through tenth sheets, corresponding to sheets one-four. These sheets contain a superset of metric values used for all figures shown. The eleventh and twelfth sheets contain batch $R^2$ and iLISI values, respectively, for the each dimension of the baseline 100-dimensional scRNA-seq embeddings used. Finally, the thirteenth sheet contains batch $R^2$ values for each dimension of the baseline 100-dimensional scATAC-seq embeddings used. For Harmony, PCA, SCA, and Scanorama, the batch $R^2$ and iLISI values reflect the embedding used for baseline benchmarking even though they were rerun for each BatchRefiner-modified approach.

### Supplementary Tables and Figures

**Supplementary Table S1.** BatchRefiner modes and approaches benchmarked for scRNA-seq. Y means the combination was benchmarked and blank means the combination was not benchmarked.

| | **All Nine Baseline Methods**[1] | | **Five Baseline Methods**[2] |
|---|---|---|---|
| **Approach** | *50 dimensions* | *100 dimensions* | *200 dimensions* |
| Scaling w/ batch $R^2$ | Y | Y | Y |
| Scaling w/ iLISI | Y | Y | Y |
| Centering w/ batch $R^2$ | | Y | |
| Centering w/ iLISI | | Y | |
| Filtering (1/5) w/ batch $R^2$ | | Y | |
| Filtering (1/5) w/ iLISI | | Y | |
| Filtering (1/3) w/ batch $R^2$ | Y | Y | Y |
| Filtering (1/3) w/ iLISI | Y | Y | Y |
| Filtering (1/2) w/ batch $R^2$ | Y | Y | Y |
| Filtering (1/2) w/ iLISI | | Y | Y |
| Filtering (batch $R^2 < 0.01$)* | | Y | |
| Filtering (batch $R^2 < 0.1$) | | Y | |
| Filtering (batch $R^2 < Q3 + IQR$) | | Y | |

[1] Concord, Harmony, LIGER, NMF, PCA, SCA, Scanorama, scVI, and Seurat CCA
[2] Concord, Harmony, PCA, SCA, and Scanorama
* Up to 1/2 (50 of 100) dimensions removed (Methods)

**Supplementary Table S2.** $P$-values for the increase in batch-correction Scores from selected BatchRefiner modes on scRNA-seq data.

| | **Scaling w/ batch $R^2$** | | **Centering w/ batch $R^2$** | |
|---|---|---|---|---|
| **Baseline Method** | T-test $p$-value | Bonferroni $p$-value | T-test $p$-value | Bonferroni $p$-value |
| Concord | 0.0146 | 0.263 | $6.82 \times 10^{-3}$ | 0.123 |
| Harmony | $1.61 \times 10^{-4}$ | $2.90 \times 10^{-3}$ | $8.98 \times 10^{-4}$ | 0.0162 |
| LIGER | $6.23 \times 10^{-3}$ | 0.112 | 0.0397 | 0.714 |
| NMF | $2.78 \times 10^{-5}$ | $5.00 \times 10^{-4}$ | $4.55 \times 10^{-6}$ | $8.18 \times 10^{-5}$ |
| PCA | $1.23 \times 10^{-5}$ | $2.21 \times 10^{-4}$ | $3.62 \times 10^{-6}$ | $6.51 \times 10^{-5}$ |
| SCA | $8.93 \times 10^{-6}$ | $1.61 \times 10^{-4}$ | $1.51 \times 10^{-8}$ | $2.71 \times 10^{-7}$ |
| Scanorama | $3.23 \times 10^{-7}$ | $5.81 \times 10^{-6}$ | $9.72 \times 10^{-5}$ | $1.75 \times 10^{-3}$ |
| scVI | $1.76 \times 10^{-7}$ | $3.17 \times 10^{-6}$ | $7.58 \times 10^{-7}$ | $1.63 \times 10^{-5}$ |
| Seurat CCA | $5.96 \times 10^{-5}$ | $1.07 \times 10^{-3}$ | 0.448 | 1 |

**Supplementary Table S3.** Dimensions remaining after BatchRefiner variable filtering approaches.

| **Baseline Method** | **Dataset**[1] | **Filtering Approach** batch $R^2 < 0.01^*$ | batch $R^2 < 0.1$ | batch $R^2 < (Q3 + IQR)$ |
|---|---|---|---|---|
| **Concord** | C elegans | 87 | 100 | 87 |
| | DKD | 50 | 100 | 95 |
| | GTEx v9 | 50 | 100 | 92 |
| | HypoMap | 69 | 100 | 87 |
| | ICA | 58 | 100 | 96 |
| | MPA | 50 | 89 | 93 |
| | Ocular Atlas | 50 | 99 | 90 |
| | Tabula Muris | 50 | 98 | 90 |
| | Tabula Sapiens | 50 | 100 | 93 |
| **Harmony** | C elegans | 94 | 100 | 83 |
| | DKD | 96 | 100 | 83 |
| | GTEx v9 | 89 | 100 | 89 |
| | HypoMap | 99 | 100 | 89 |
| | ICA | 92 | 99 | 84 |
| | MPA | 52 | 92 | 89 |
| | Ocular Atlas | 64 | 96 | 86 |
| | Tabula Muris | 75 | 97 | 82 |
| | Tabula Sapiens | 88 | 100 | 90 |
| **LIGER** | C elegans | 81 | 100 | 91 |
| | DKD | 84 | 99 | 90 |
| | GTEx v9[†] | 20 | 37 | 39 |
| | HypoMap | 94 | 100 | 87 |
| | ICA | 83 | 100 | 87 |
| | MPA | 33 | 92 | 91 |
| | Ocular Atlas | 50 | 94 | 85 |
| | Tabula Muris | 50 | 96 | 84 |
| | Tabula Sapiens[‡] | 42 | 82 | 76 |
| **NMF** | C elegans | 50 | 92 | 92 |
| | DKD | 50 | 84 | 89 |
| | GTEx v9 | 50 | 81 | 90 |
| | HypoMap | 64 | 98 | 87 |
| | ICA | 50 | 78 | 89 |
| | MPA | 50 | 28 | 93 |
| | Ocular Atlas | 50 | 83 | 84 |
| | Tabula Muris | 50 | 80 | 91 |
| | Tabula Sapiens | 50 | 84 | 88 |

[1] ICA, Immune Cell Atlas; MPA, Mouse Pancreas Atlas
[*] Up to half (50) of dimensions removed, *i.e,* the batch $R^2$ threshold is the greater of 0.01 and the median value.
[†] 40 initial dimensions
[‡] 83 initial dimensions

*Continued on next page*

| **Method** | **Dataset**[1] | batch $R^2 < 0.01^*$ | batch $R^2 < 0.1$ | batch $R^2 < (Q3 + IQR)$ |
|---|---|---|---|---|
| **PCA** | C elegans | 50 | 96 | 90 |
| | DKD | 50 | 86 | 88 |
| | GTEx v9 | 50 | 88 | 90 |
| | HypoMap | 77 | 99 | 90 |
| | ICA | 50 | 89 | 89 |
| | MPA | 50 | 70 | 92 |
| | Ocular Atlas | 50 | 85 | 88 |
| | Tabula Muris | 50 | 93 | 82 |
| | Tabula Sapiens | 50 | 75 | 96 |
| **SCA** | C elegans | 50 | 95 | 86 |
| | DKD | 50 | 82 | 90 |
| | GTEx v9 | 50 | 85 | 90 |
| | HypoMap | 55 | 100 | 96 |
| | ICA | 50 | 83 | 89 |
| | MPA | 50 | 21 | 89 |
| | Ocular Atlas | 50 | 81 | 90 |
| | Tabula Muris | 50 | 86 | 93 |
| | Tabula Sapiens | 50 | 63 | 97 |
| **Scanorama** | C elegans | 83 | 99 | 87 |
| | DKD | 50 | 96 | 93 |
| | GTEx v9 | 50 | 87 | 92 |
| | HypoMap | 66 | 100 | 92 |
| | ICA | 50 | 98 | 90 |
| | MPA | 50 | 61 | 95 |
| | Ocular Atlas | 50 | 91 | 88 |
| | Tabula Muris | 50 | 92 | 87 |
| | Tabula Sapiens | 50 | 74 | 94 |
| **scVI** | C elegans | 50 | 99 | 89 |
| | DKD | 50 | 99 | 90 |
| | GTEx v9 | 50 | 98 | 85 |
| | HypoMap | 63 | 100 | 83 |
| | ICA | 50 | 99 | 93 |
| | MPA | 50 | 66 | 84 |
| | Ocular Atlas | 50 | 90 | 93 |
| | Tabula Muris | 50 | 100 | 95 |
| | Tabula Sapiens | 50 | 96 | 84 |
| **Seurat CCA** | C elegans | 89 | 99 | 83 |
| | DKD | 82 | 100 | 90 |
| | GTEx v9 | 79 | 100 | 85 |
| | HypoMap | 99 | 100 | 83 |
| | ICA | 77 | 100 | 93 |
| | MPA | 58 | 100 | 84 |
| | Ocular Atlas | 85 | 100 | 85 |
| | Tabula Muris | 53 | 98 | 89 |
| | Tabula Sapiens | 80 | 100 | 84 |

[1] ICA, Immune Cell Atlas; MPA, Mouse Pancreas Atlas

* Up to half (50) columns removed, *i.e,* the batch $R^2$ threshold is the greater of 0.01 and the median value.

**Supplementary Table S4.** $P$-values for the increase in batch-correction scores from selected BatchRefiner modes on scATAC-seq data.

| | **Scaling w/ batch $R^2$** | | **Centering w/ batch $R^2$** | |
|---|---|---|---|---|
| **Baseline Method** | T-test $p$-value | Bonferroni $p$-value | T-test $p$-value | Bonferroni $p$-value |
| pycisTopic | $6.58 \times 10^{-5}$ | $1.32 \times 10^{-3}$ | $4.21 \times 10^{-7}$ | $8.41 \times 10^{-6}$ |
| pycisTopic+Harmony | $2.43 \times 10^{-3}$ | 0.0486 | $1.10 \times 10^{-3}$ | 0.0221 |
| peakVI | $8.82 \times 10^{-3}$ | 0.176 | $2.46 \times 10^{-6}$ | $4.92 \times 10^{-5}$ |
| Signac | $6.30 \times 10^{-6}$ | $1.26 \times 10^{-4}$ | $8.55 \times 10^{-5}$ | $1.71 \times 10^{-3}$ |
| Signac+Harmony | 0.0161 | 0.322 | 0.758 | 1 |
| Signac+Seurat | $6.00 \times 10^{-3}$ | 0.120 | 0.0162 | 0.323 |
| SnapATAC2 | $4.78 \times 10^{-6}$ | $9.57 \times 10^{-5}$ | $9.99 \times 10^{-9}$ | $2.00 \times 10^{-7}$ |
| SnapATAC2+Harmony | $1.43 \times 10^{-3}$ | 0.0291 | 0.0838 | 1 |
| SVD | $1.43 \times 10^{-6}$ | $3.25 \times 10^{-5}$ | $8.84 \times 10^{-8}$ | $1.77 \times 10^{-6}$ |
| SVD+Harmony | $2.05 \times 10^{-4}$ | $4.10 \times 10^{-3}$ | $8.76 \times 10^{-5}$ | $1.75 \times 10^{-3}$ |

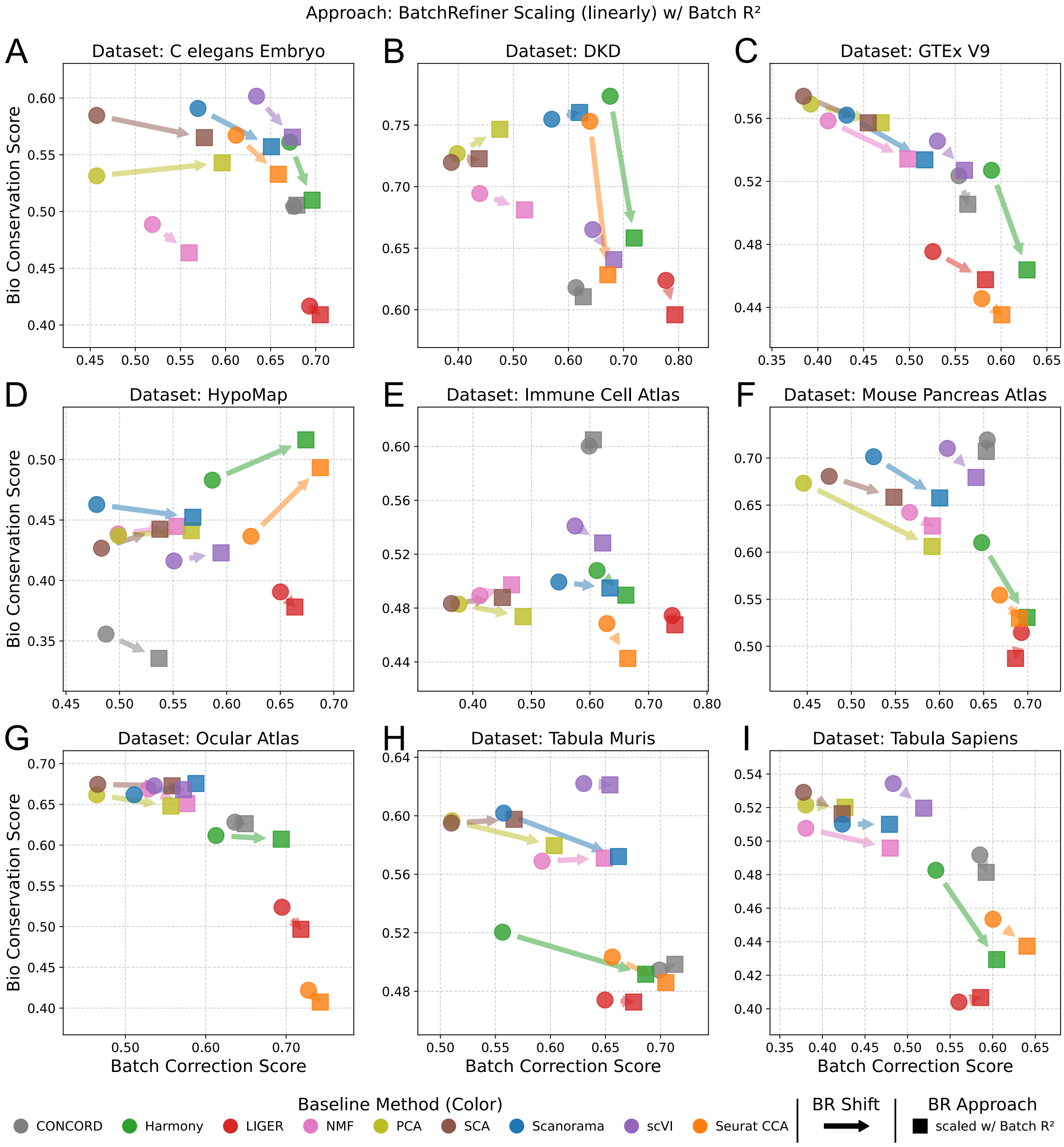


**Supplementary Fig. S1.** Benchmarks of BatchRefiner scaled embeddings on individual scRNA-seq datasets. Benchmarking of baseline and BatchRefiner-**scaled** embeddings, shown as mean bio-conservation (y) and batch-integration (x) scores for each of nine scRNA-seq datasets individually. BatchRefiner-scaled embeddings using batch $R^2$ as the dimension-evaluation metric are shown as squares. Baseline 100-dimensional embeddings from nine methods are shown as circles, colored by method as shown in the bottom left. Shifts from baseline methods to the corresponding scaled embedding are shown with arrows.

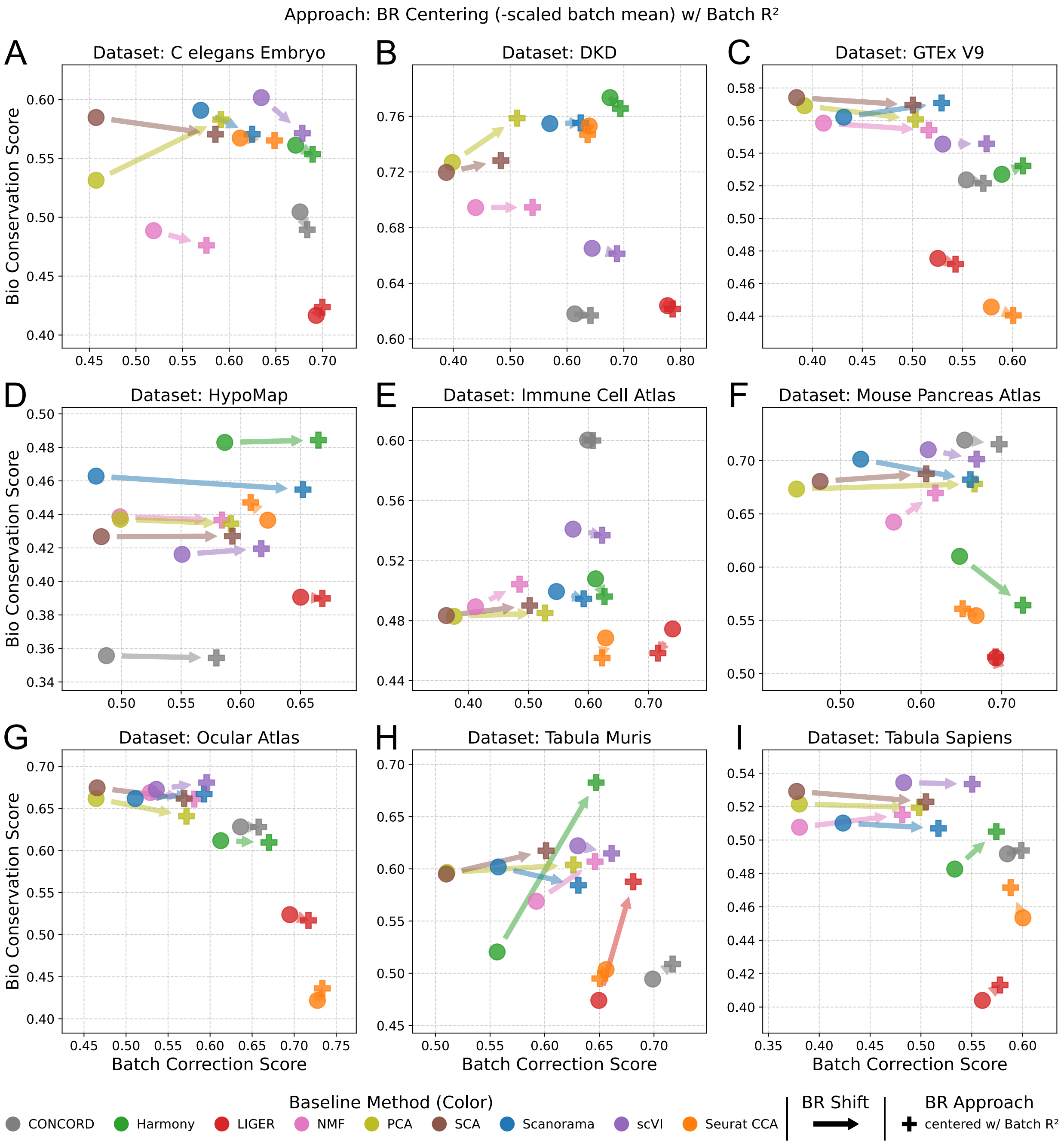


**Supplementary Fig. S2.** Benchmarks of BatchRefiner centered embeddings on individual scRNA-seq datasets. Benchmarking of baseline and BatchRefiner-**centered** embeddings, shown as mean bio-conservation (y) and batch-integration (x) scores for each of nine scRNA-seq datasets individually. BatchRefiner-centered embeddings using batch $R^2$ as the dimension-evaluation metric are shown as plus symbols. Baseline 100-dimensional embeddings from nine methods are shown as circles, colored by method as shown in the bottom left. Shifts from baseline methods to the corresponding centered embedding are shown with arrows.

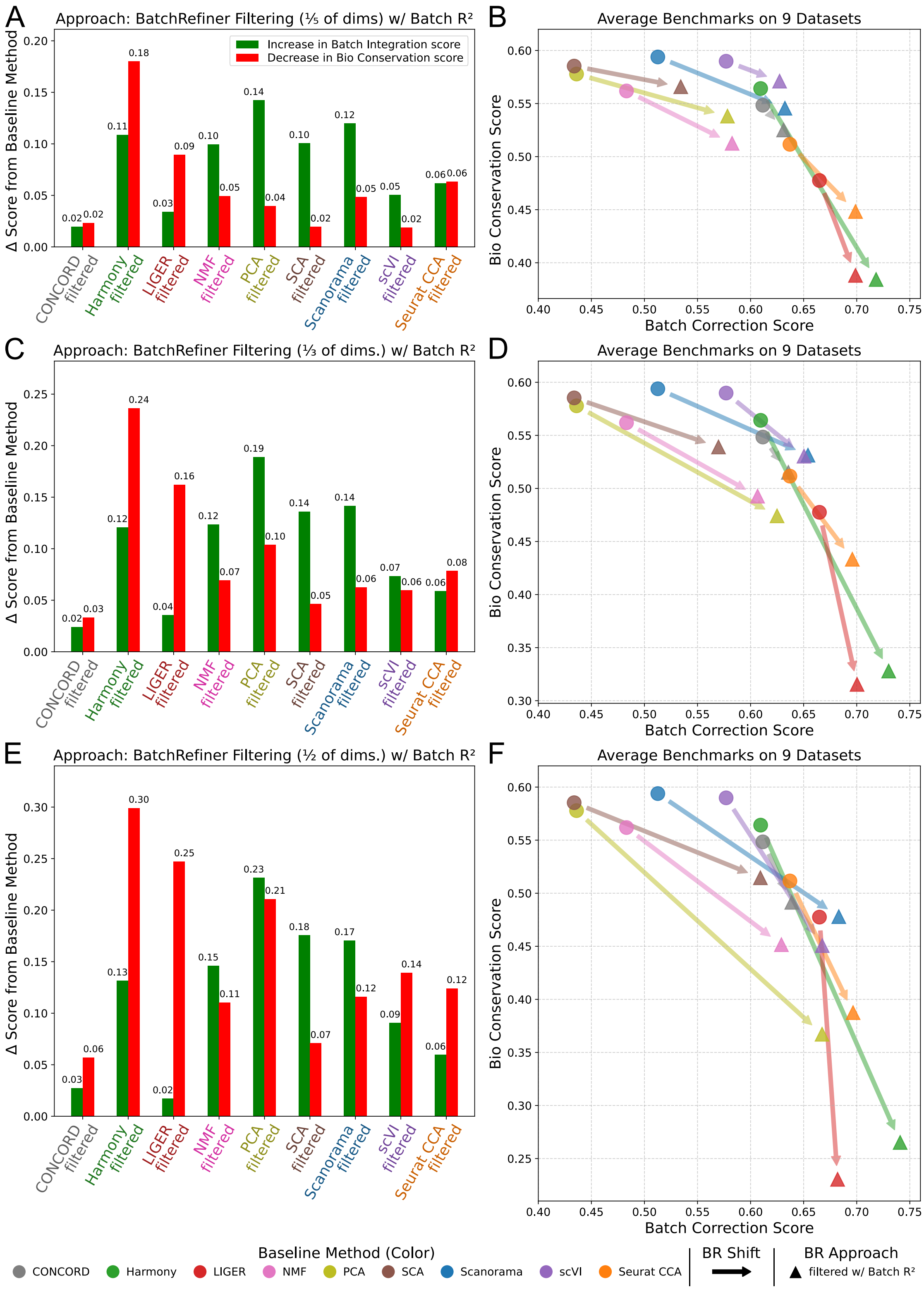


**Supplementary Fig. S3.** Benchmarking BatchRefiner **filtering**. **(A)** Shifts in average batch-integration score (increases, green) and average bio-conservation score (decreases, red) relative to the corresponding baseline method for BatchRefiner filtering using batch $R^2$ to remove the lowest-scoring **fifth** of embedding dimensions. Shifts are averaged across nine scRNA-seq datasets. **(B)** Mean bio-conservation (y) and batch integration (x) scores of baseline and BatchRefiner-filtered embeddings. BatchRefiner-filtered embeddings as in (A) are shown as triangles. Baseline 100-dimensional embeddings from ten methods are shown as circles, colored by method as shown in the bottom left. Shifts from baseline methods to the corresponding filtered embedding are shown with arrows. **(C-D)** Benchmarking **filtering** the lowest-scoring **third** of dimensions using batch $R^2$. Data are shown as in (A-B). **(E-F)** Benchmarking **filtering** the lowest-scoring **half** of dimensions using batch $R^2$. Data are shown as in (A-B).

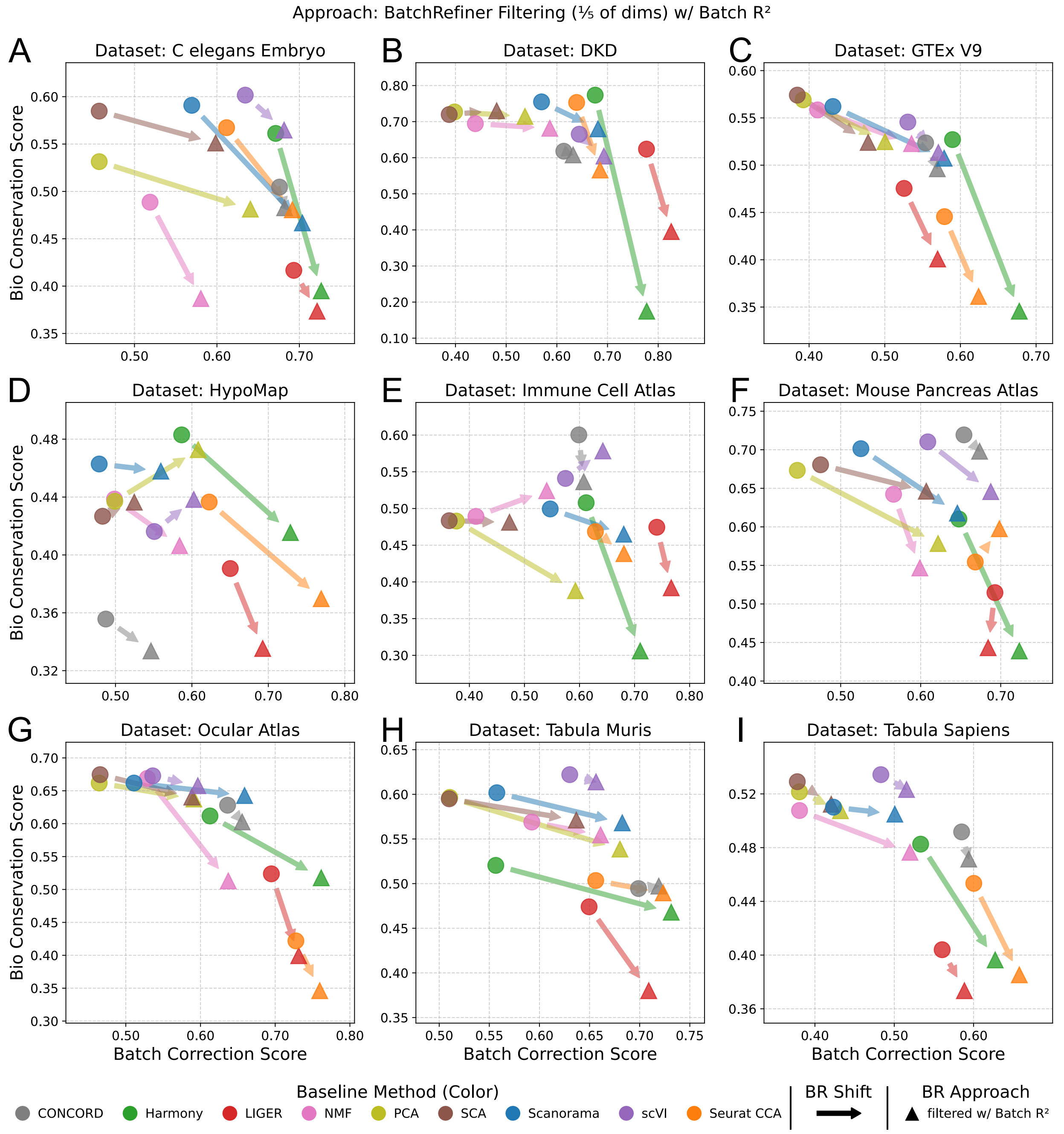


**Supplementary Fig. S4.** Benchmarking BatchRefiner filtering one fifth of dimensions on individual scRNA-seq datasets. Benchmarking of baseline and BatchRefiner-**filtered** embeddings, shown as mean bio-conservation (y) and batch-integration (x) scores for each of nine scRNA-seq datasets individually. BatchRefiner-filtered embeddings using batch $R^2$ as the dimension-evaluation metric, to filter one **fifth** of dimensions, are shown as triangles. Baseline 100-dimensional embeddings from nine methods are shown as circles, colored by method as shown in the bottom left. Shifts from baseline methods to the corresponding filtered embedding are shown with arrows.

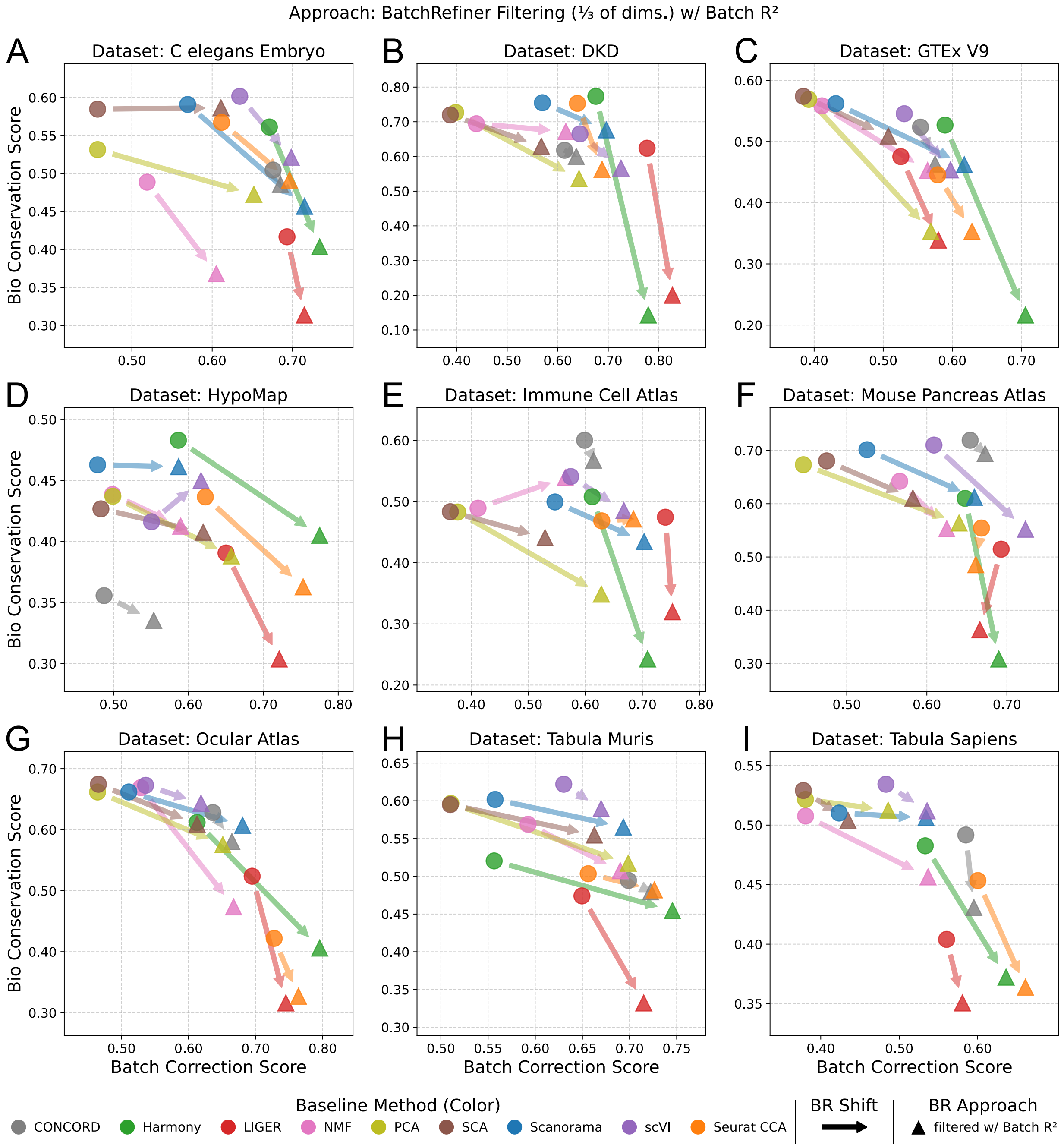


**Supplementary Fig. S5.** Benchmarking BatchRefiner filtering one third of dimensions on individual scRNA-seq datasets. Benchmarking of baseline and BatchRefiner-**filtered** embeddings, shown as mean bio-conservation (y) and batch-integration (x) scores for each of nine scRNA-seq datasets individually. BatchRefiner-filtered embeddings using batch $R^2$ as the dimension-evaluation metric, to filter one **third** of dimensions, are shown as triangles. Baseline 100-dimensional embeddings from nine methods are shown as circles, colored by method as shown in the bottom left. Shifts from baseline methods to the corresponding filtered embedding are shown with arrows.

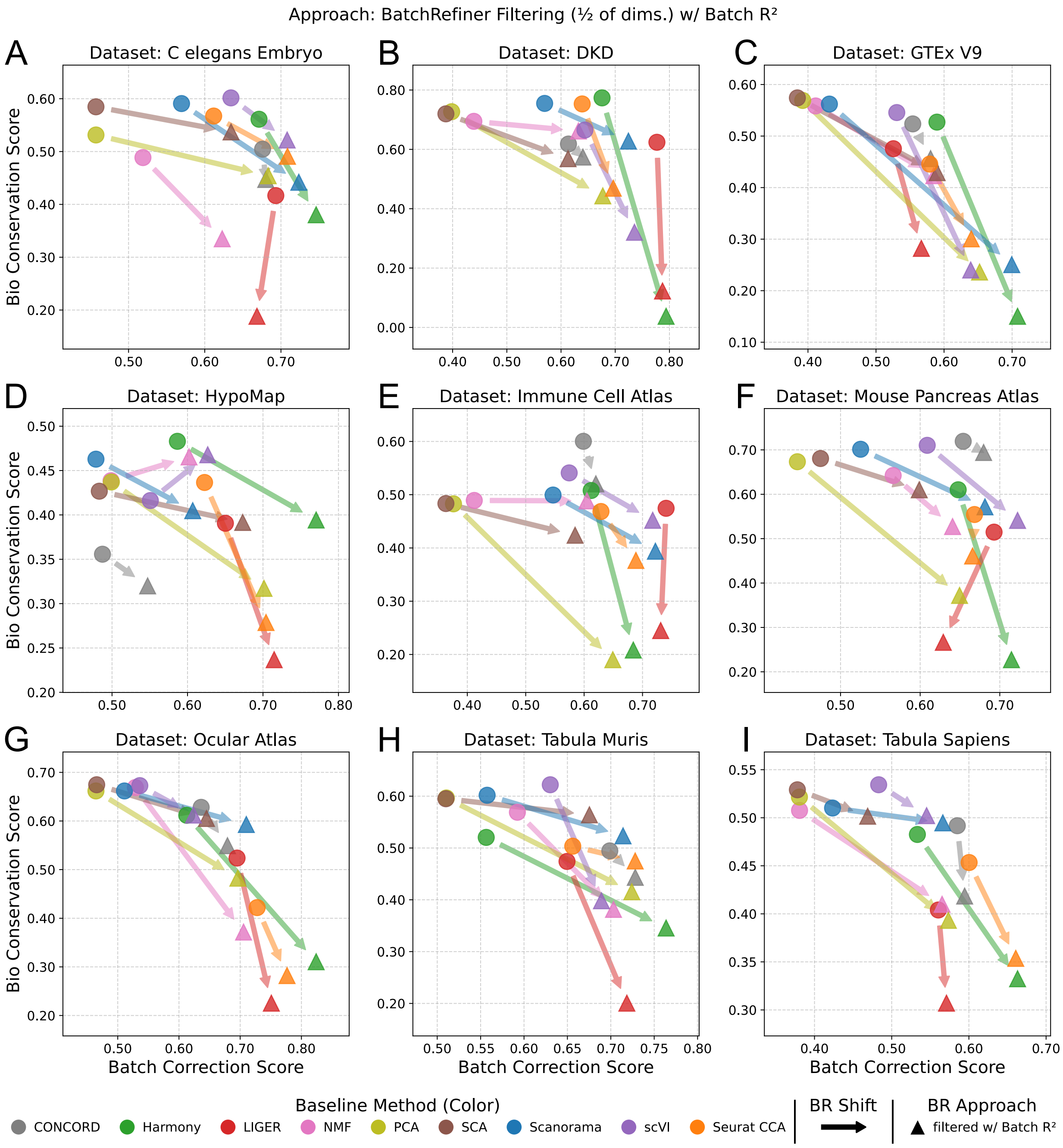


**Supplementary Fig. S6.** Benchmarking BatchRefiner filtering one half of dimensions on individual scRNA-seq datasets. Benchmarking of baseline and BatchRefiner-**filtered** embeddings, shown as mean bio-conservation (y) and batch-integration (x) scores for each of nine scRNA-seq datasets individually. BatchRefiner-filtered embeddings using batch $R^2$ as the dimension-evaluation metric, to filter one **half** of dimensions, are shown as triangles. Baseline 100-dimensional embeddings from nine methods are shown as circles, colored by method as shown in the bottom left. Shifts from baseline methods to the corresponding filtered embedding are shown with arrows.

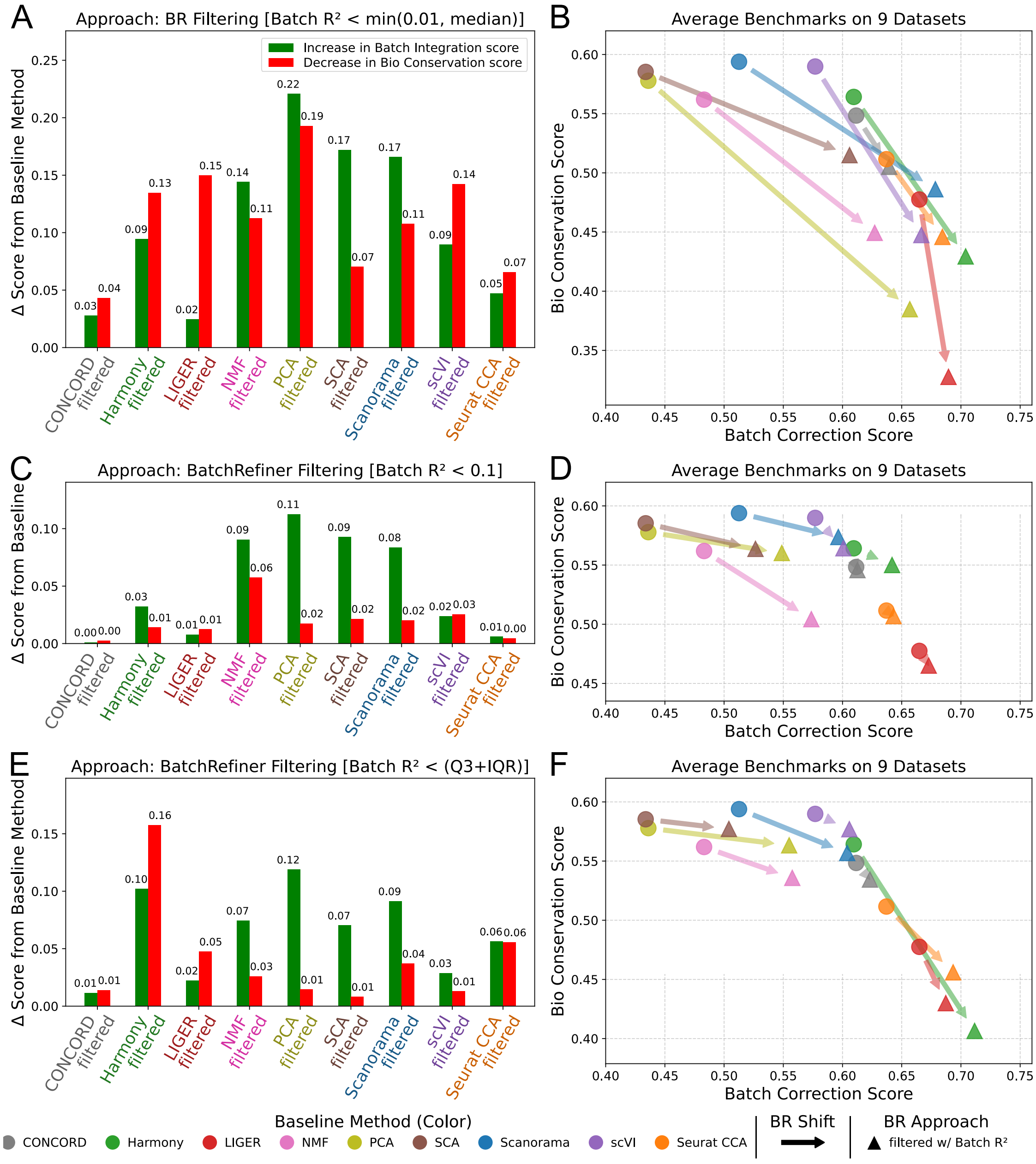


**Supplementary Fig. S7.** Benchmarking additional BatchRefiner **filtering** approaches. **(A)** Shifts in average batch-integration score (increases, green) and average bio-conservation score (decreases, red) relative to the corresponding baseline method for BatchRefiner filtering, removing dimensions with batch $R^2 \geq 0.01$, up to half of the columns. Shifts are averaged across nine scRNA-seq datasets. **(B)** Benchmarking of baseline and BatchRefiner-filtered embeddings of OpenProblems datasets, shown as mean bio-conservation (y) and batch-integration (x) scores, averaged across datasets. BatchRefiner-filtered embeddings (removing dimensions where $R^2 \geq 0.01$, up to 50%) as in (A) are shown as triangles. Baseline 100-dimensional embeddings from nine methods are shown as circles, colored by method as shown in the bottom left. Shifts from baseline methods to the corresponding scaled embedding are shown with arrows. **(C-D)** Benchmarking BatchRefiner filtering, removing only dimensions with batch $R^2 \geq 0.1$. Data are shown as in (A-B). **(E-F)** Benchmarking BatchRefiner filtering, instead removing dimensions with batch $R^2$ greater than the third quartile plus the inter-quartile range of $R^2$ scores. Data are shown as in (A-B).

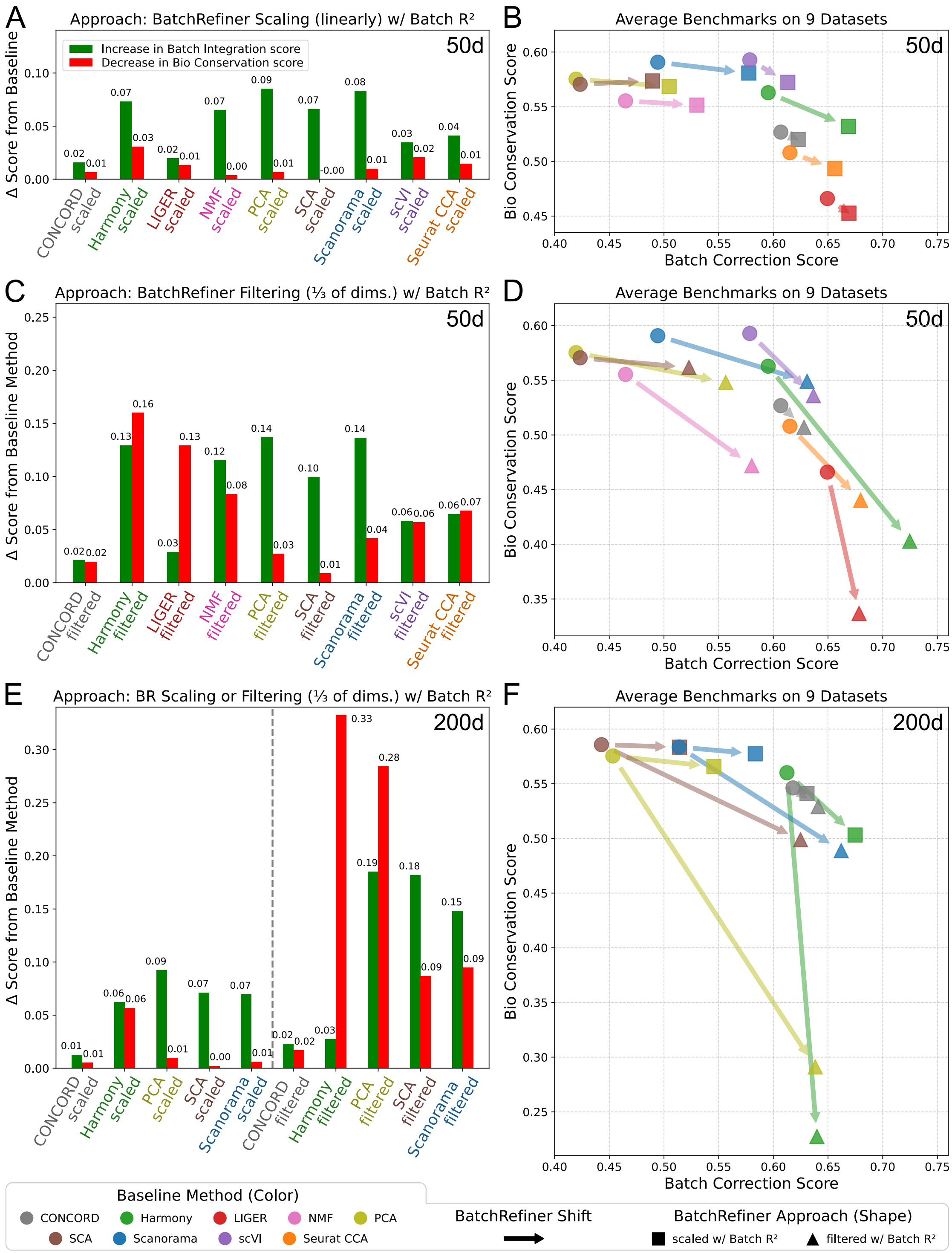


**Supplementary Fig. S8.** Benchmarking BatchRefiner with varied initial embedding dimension. **(A)** Shifts in average batch-integration score (increases, green) and average bio-conservation score (decreases, red) relative to the corresponding baseline method for BatchRefiner **scaling** of **50-dimension** embeddings using batch $R^2$. Shifts are averaged across nine scRNA-seq datasets. **(B)** Benchmarking of baseline and BatchRefiner-scaled 50d embeddings of nine scRNA-seq datasets, shown as mean bio-conservation (y) and batch-integration (x) scores, averaged across datasets. BatchRefiner-scaled embeddings are shown as squares. Baseline 50-dimension embeddings from nine methods are shown as circles, colored by method (bottom left). Shifts from baseline methods to the corresponding scaled embedding are shown with arrows. **(C-D)** Benchmarking BatchRefiner **filtering**, removing the lowest-scoring third of 50 initial dimensions using batch $R^2$. Data are shown as in (A-B), except that filtered embeddings are shown as in triangles. **(E-F)** Benchmarking BatchRefiner **scaling and filtering** (one third of dimensions), using batch $R^2$, of **200-dimension** embeddings from five baseline methods. Data are shown as in (A-D).

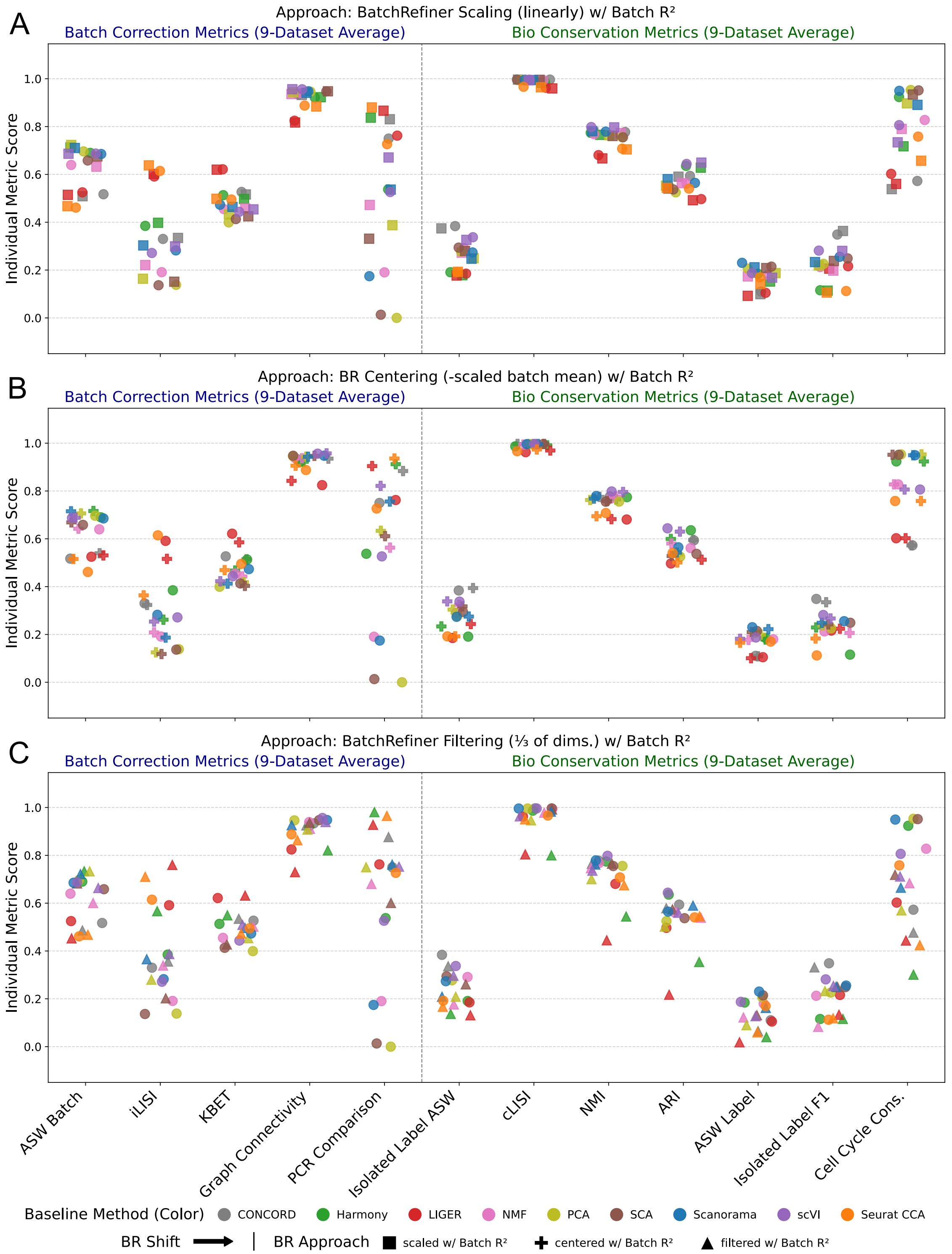


**Supplementary Fig. S9.** Individual OpenProblems benchmarks for baseline scRNA-seq embeddings and BatchRefiner-modified embeddings using batch $R^2$. **(A)** Values of each of 12 individual metrics (batch-integration metrics on left, bio-conservation metrics on right) for each of nine baseline and corresponding BatchRefiner-scaled embeddings, averaged across nine scRNA-seq datasets datasets. Baseline 100-dimension embeddings are shown as circles, colored by method (bottom). BatchRefiner-scaled embeddings are shown as squares. **(B)** Individual metric values for baseline and corresponding BatchRefiner-centered embeddings. Data are shown as in (A); BatchRefiner-centered embeddings are shown as plus symbols. **(C)** Individual metric values for baseline and corresponding BatchRefiner-filtered (one-**third** of dimensions) embeddings. Data are shown as in (A); BatchRefiner-filtered embeddings are shown as triangles.

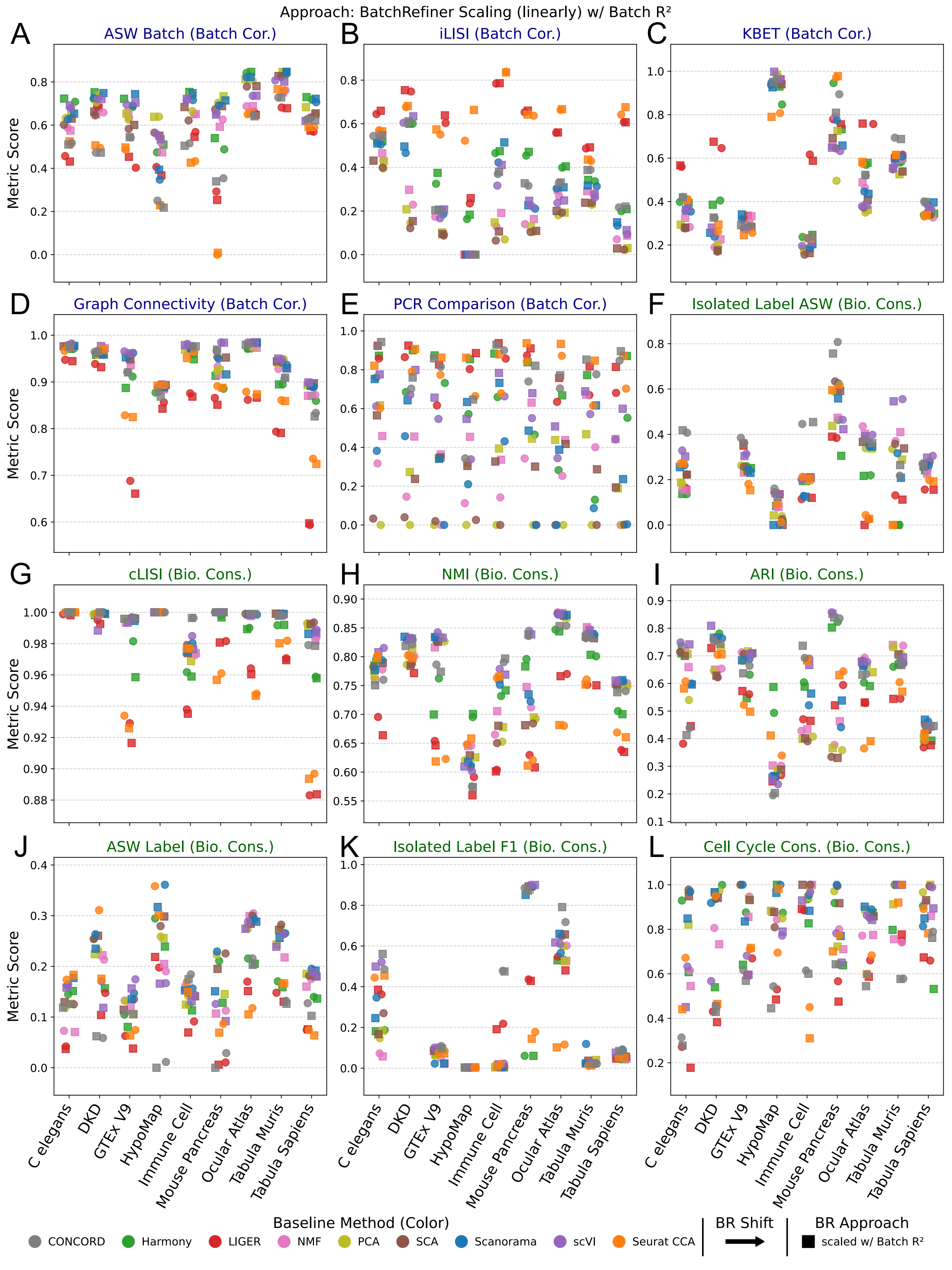


**Supplementary Fig. S10.** Per-dataset OpenProblems benchmarks for baseline scRNA-seq embeddings and BatchRefiner **scaled** embeddings using batch $R^2$. Values of each of 12 individual metrics (batch-integration metrics on left, bio-conservation metrics on right) for each of nine baseline and corresponding BatchRefiner-scaled embeddings, and for each of nine scRNA-seq datasets (one per panel). Baseline 100-dimension embeddings from nine methods are shown as circles, colored by method as shown in the bottom left. BatchRefiner-scaled embeddings are shown as squares.

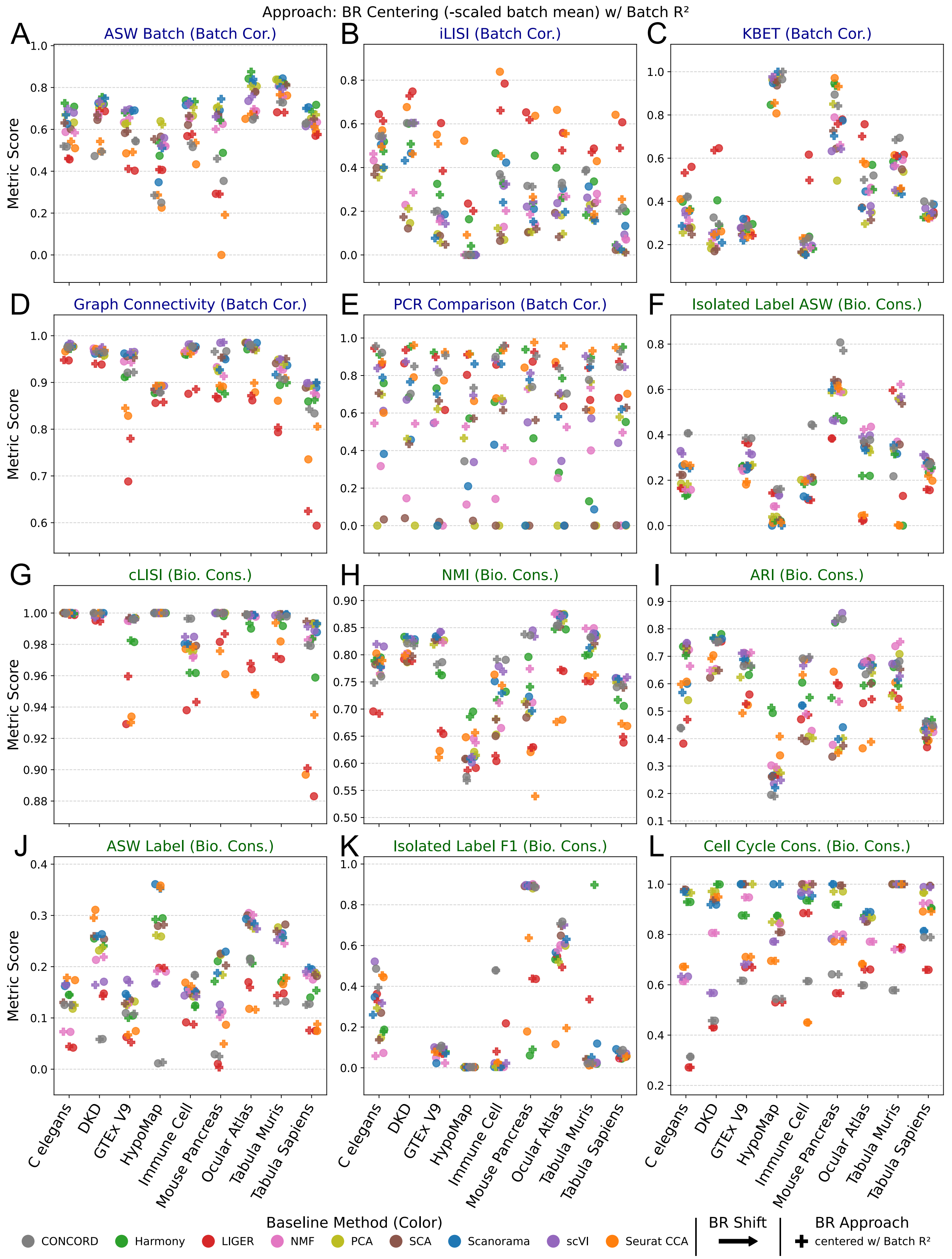


**Supplementary Fig. S11.** Per-dataset OpenProblems benchmarks for baseline scRNA-seq embeddings and BatchRefiner **centered** embeddings using batch $R^2$. Values of each of 12 individual metrics (batch-integration metrics on left, bio-conservation metrics on right) for each of nine baseline and corresponding BatchRefiner-scaled embeddings, and for each of nine scRNA-seq datasets (one per panel). Baseline 100-dimension embeddings from nine methods are shown as circles, colored by method as shown in the bottom left. BatchRefiner-centered embeddings are shown as plus symbols.

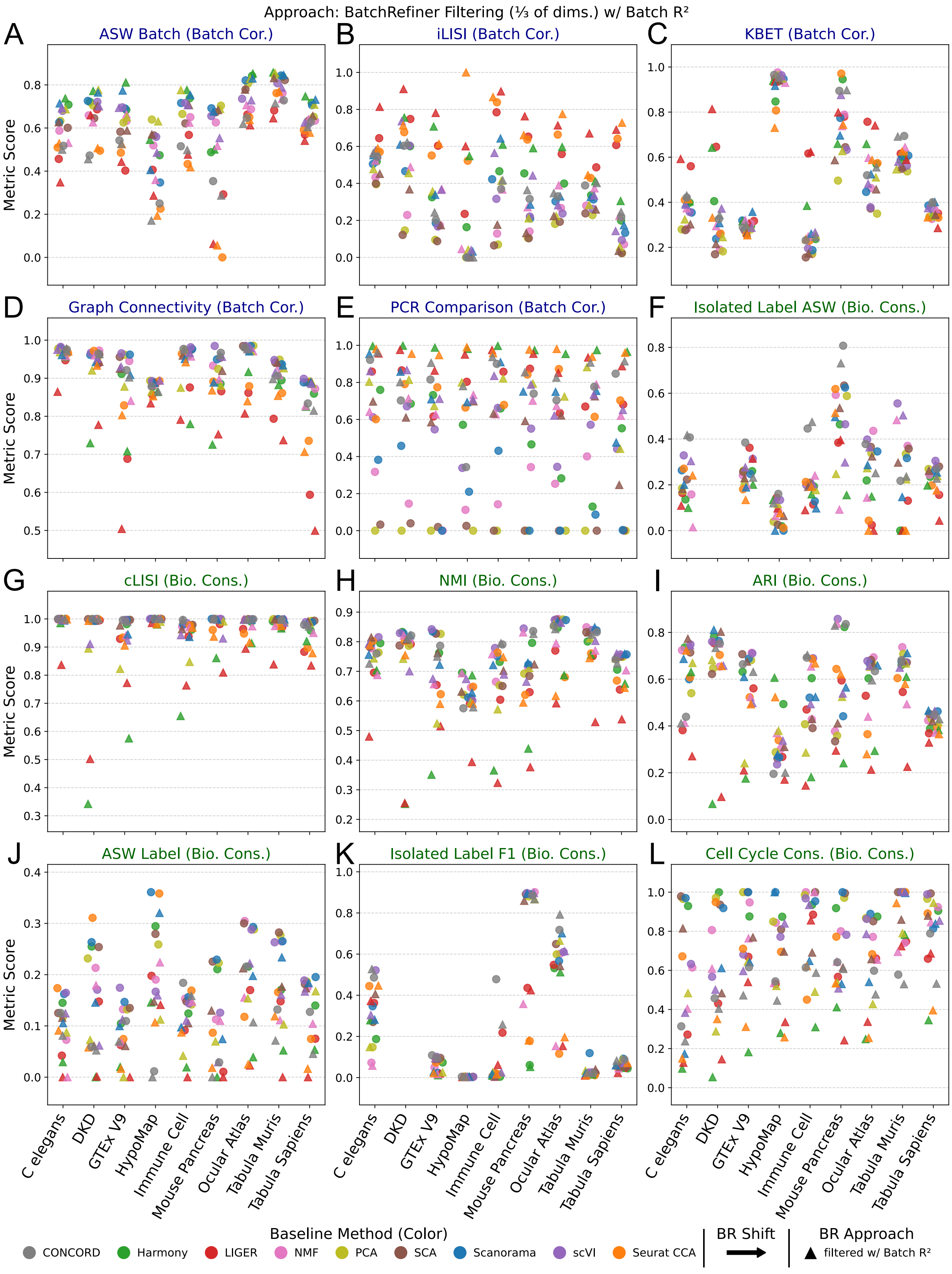


**Supplementary Fig. S12.** Per-dataset OpenProblems benchmarks for baseline scRNA-seq embeddings and BatchRefiner **filtered** embeddings using batch $R^2$. Values of each of 12 individual metrics (batch-integration metrics on left, bio-conservation metrics on right) for each of nine baseline and corresponding BatchRefiner-scaled embeddings, and for each of nine scRNA-seq datasets (one per panel). Baseline 100-dimension embeddings from nine methods are shown as circles, colored by method as shown in the bottom left. BatchRefiner-filtered embeddings (lowest-scoring **third** of dimensions removed) are shown as triangles.

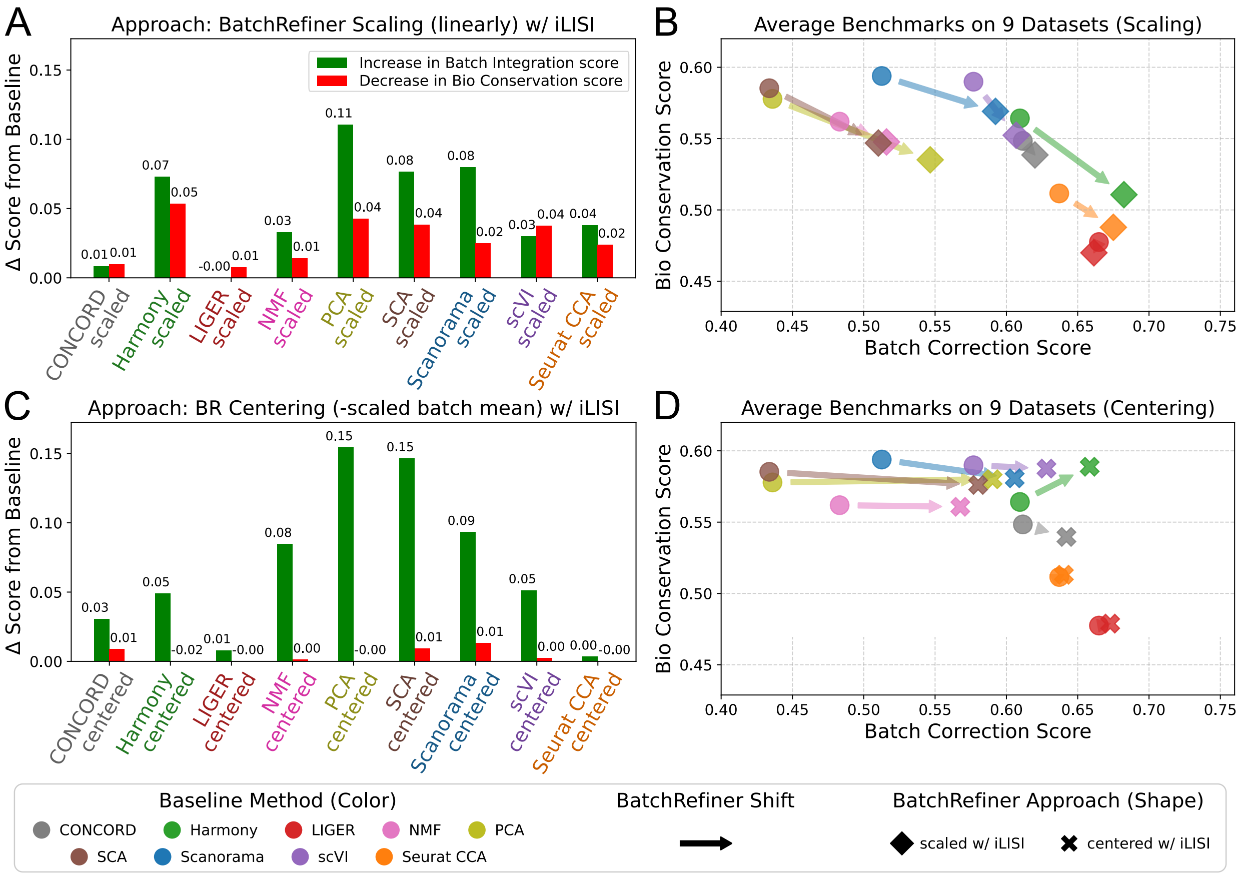


**Supplementary Fig. S13.** Benchmarking BatchRefiner **scaling and centering** using iLISI. **(A)** Shifts in average batch-integration score (increases, green) and average bio-conservation score (decreases, red) relative to the corresponding baseline method for BatchRefiner scaling using iLISI. Shifts are averaged across nine scRNA-seq datasets. **(B)** Benchmarking of baseline and BatchRefiner-scaled embeddings of OpenProblems datasets, shown as mean bio-conservation (y) and batch-integration (x) scores, averaged across datasets. BatchRefiner-scaled embeddings as in (A) are shown as triangles. Baseline 100-dimensional embeddings from nine methods are shown as circles, colored by method as shown in the bottom left. Shifts from baseline methods to the corresponding scaled embedding are shown with arrows. **(C-D)** Changes in average benchmark scores, as in (A-B), for BatchRefiner (BR) centering (subtracting the batch mean, weighted by iLISI, from each dimension; X symbols).

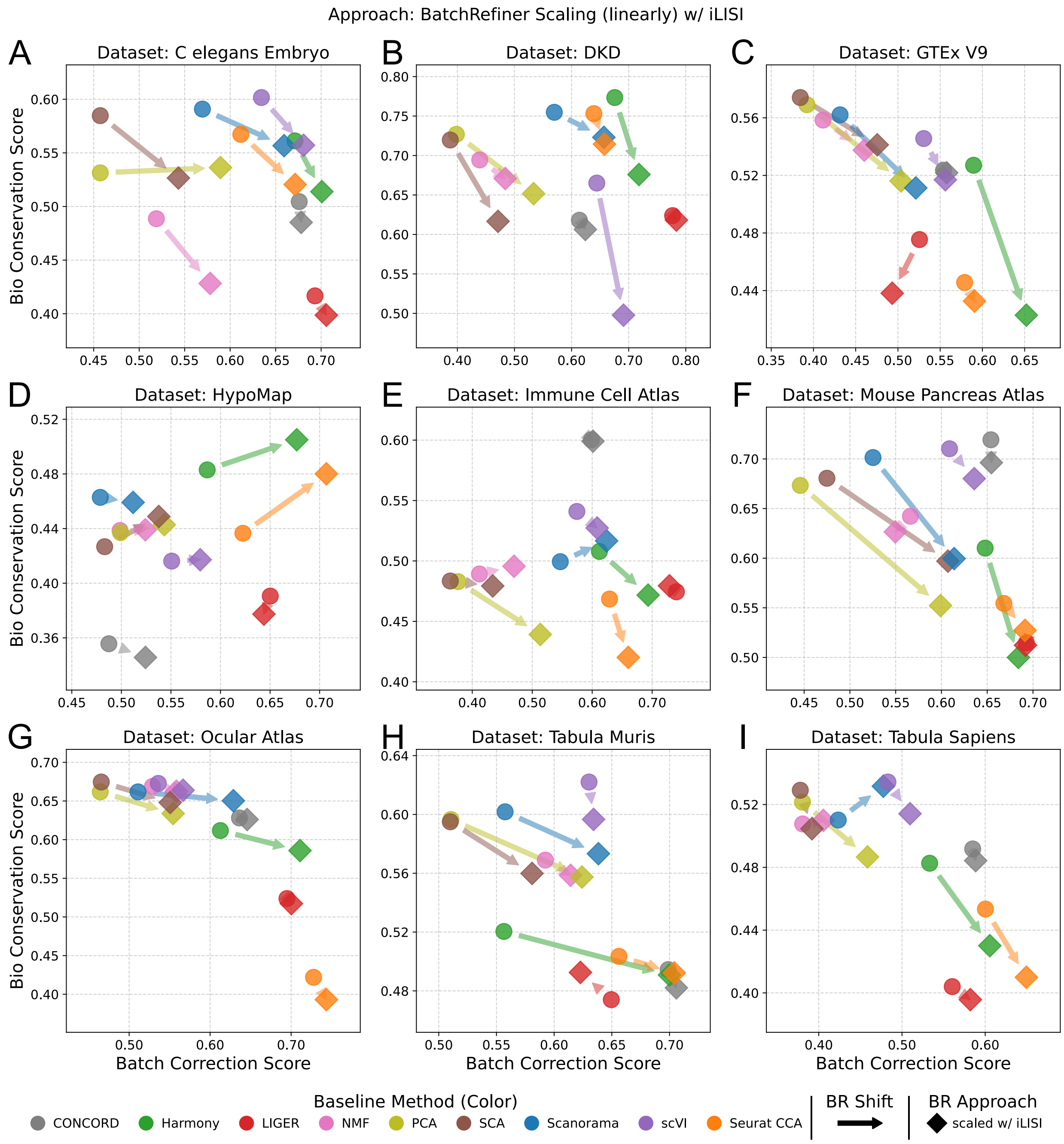


**Supplementary Fig. S14.** Benchmarks of BatchRefiner scaled embeddings, using iLISI, on individual scRNA-seq datasets. Benchmarking of baseline and BatchRefiner-**scaled** embeddings, shown as mean bio-conservation (y) and batch-integration (x) scores for each of nine scRNA-seq datasets individually. BatchRefiner-scaled embeddings using iLISI as the dimension-evaluation metric are shown as trianges. Baseline 100-dimensional embeddings from nine methods are shown as circles, colored by method as shown in the bottom left. Shifts from baseline methods to the corresponding scaled embedding are shown with arrows.

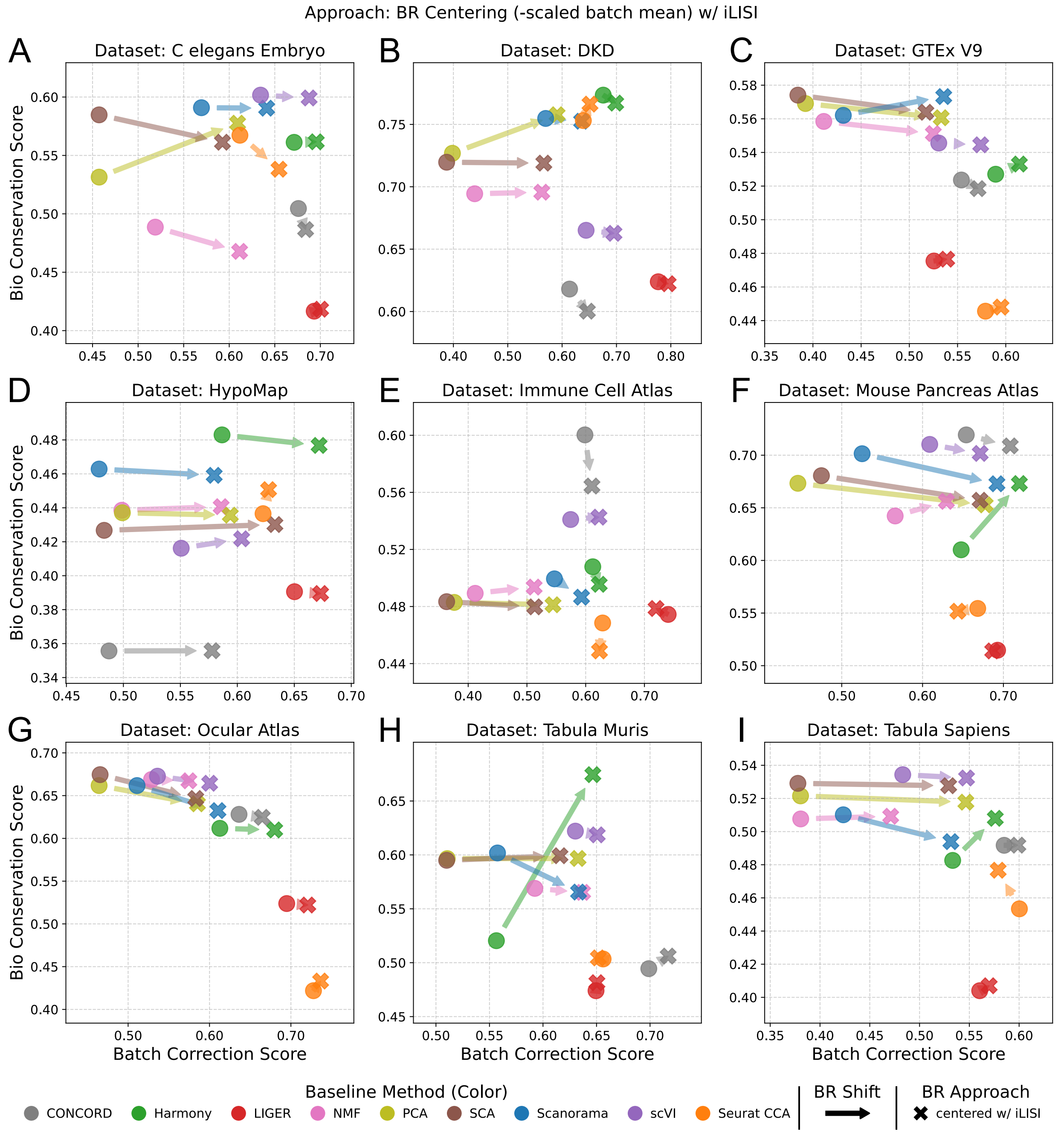


**Supplementary Fig. S15.** Benchmarks of BatchRefiner centered embeddings, using iLISI, on individual scRNA-seq datasets. Benchmarking of baseline and BatchRefiner-**centered** embeddings, shown as mean bio-conservation (y) and batch-integration (x) scores for each of nine scRNA-seq datasets individually. BatchRefiner-centered embeddings using iLISI as the dimension-evaluation metric are shown as X symbols. Baseline 100-dimensional embeddings from nine methods are shown as circles, colored by method as shown in the bottom left. Shifts from baseline methods to the corresponding centered embedding are shown with arrows.

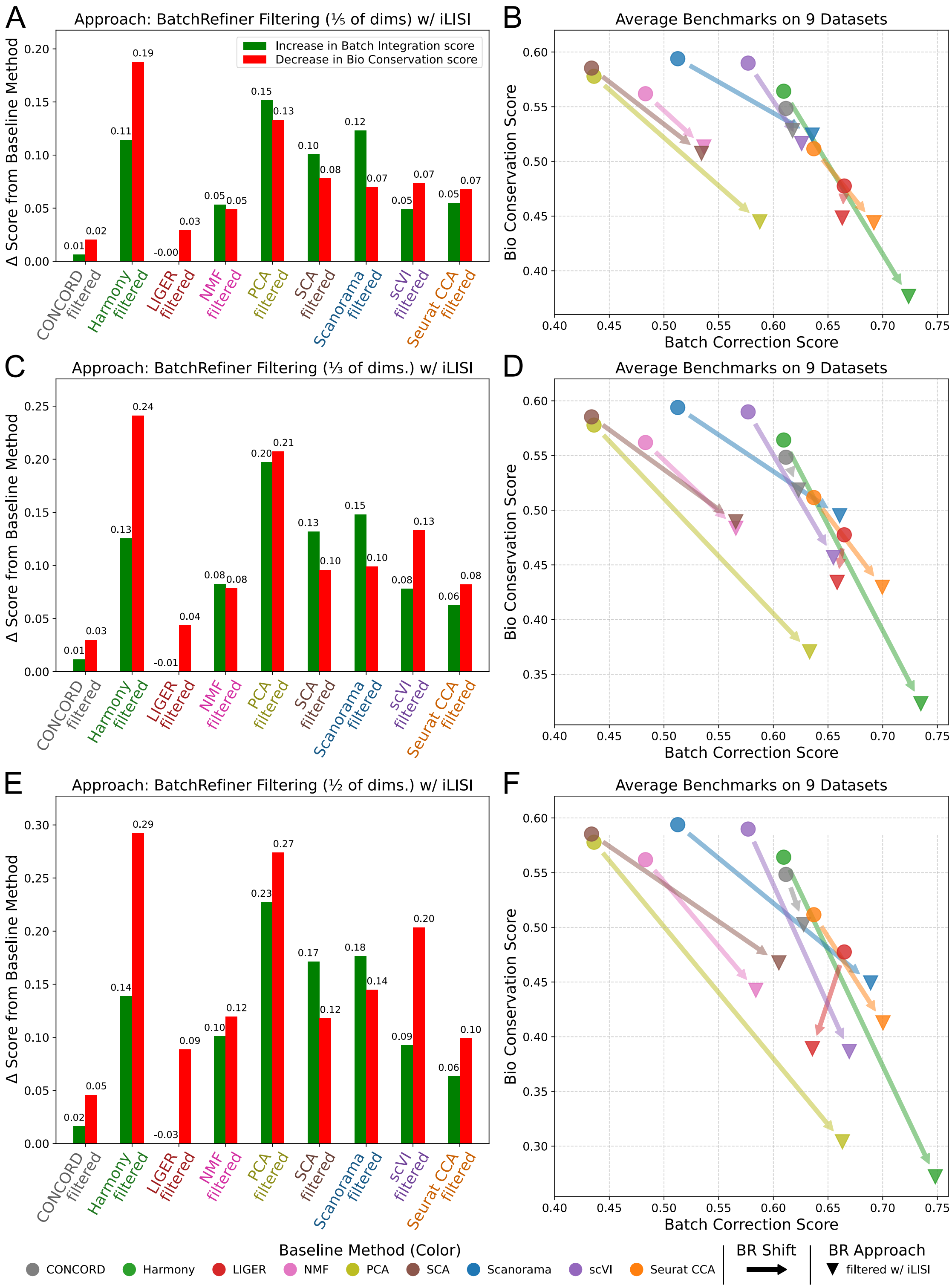


**Supplementary Fig. S16.** Benchmarking BatchRefiner **filtering** fixed fractions of dimensions using iLISI. **(A)** Shifts in average batch-integration score (increases, green) and average bio-conservation score (decreases, red) relative to the corresponding baseline method for BatchRefiner filtering using iLISI to remove the lowest-scoring **fifth** of embedding dimensions. Shifts are averaged across nine scRNA-seq datasets. **(B)** Benchmarking of baseline and BatchRefiner-filtered (1/5) embeddings of OpenProblems datasets, shown as mean bio-conservation (y) and batch-integration (x) scores, averaged across datasets. BatchRefiner-filtered embeddings as in (A) are shown as triangles. Baseline 100-dimensional embeddings from ten methods are shown as circles, colored by method as shown in the bottom left. Shifts from baseline methods to the corresponding filtered embedding are shown with arrows. **(C-D)** Benchmarking BatchRefiner filtering, removing the lowest-scoring **third** of dimensions using iLISI. Data are shown as in (A-B). **(E-F)** Benchmarking BatchRefiner filtering, removing the lowest-scoring **half** of dimensions using iLISI. Data are shown as in (A-B).

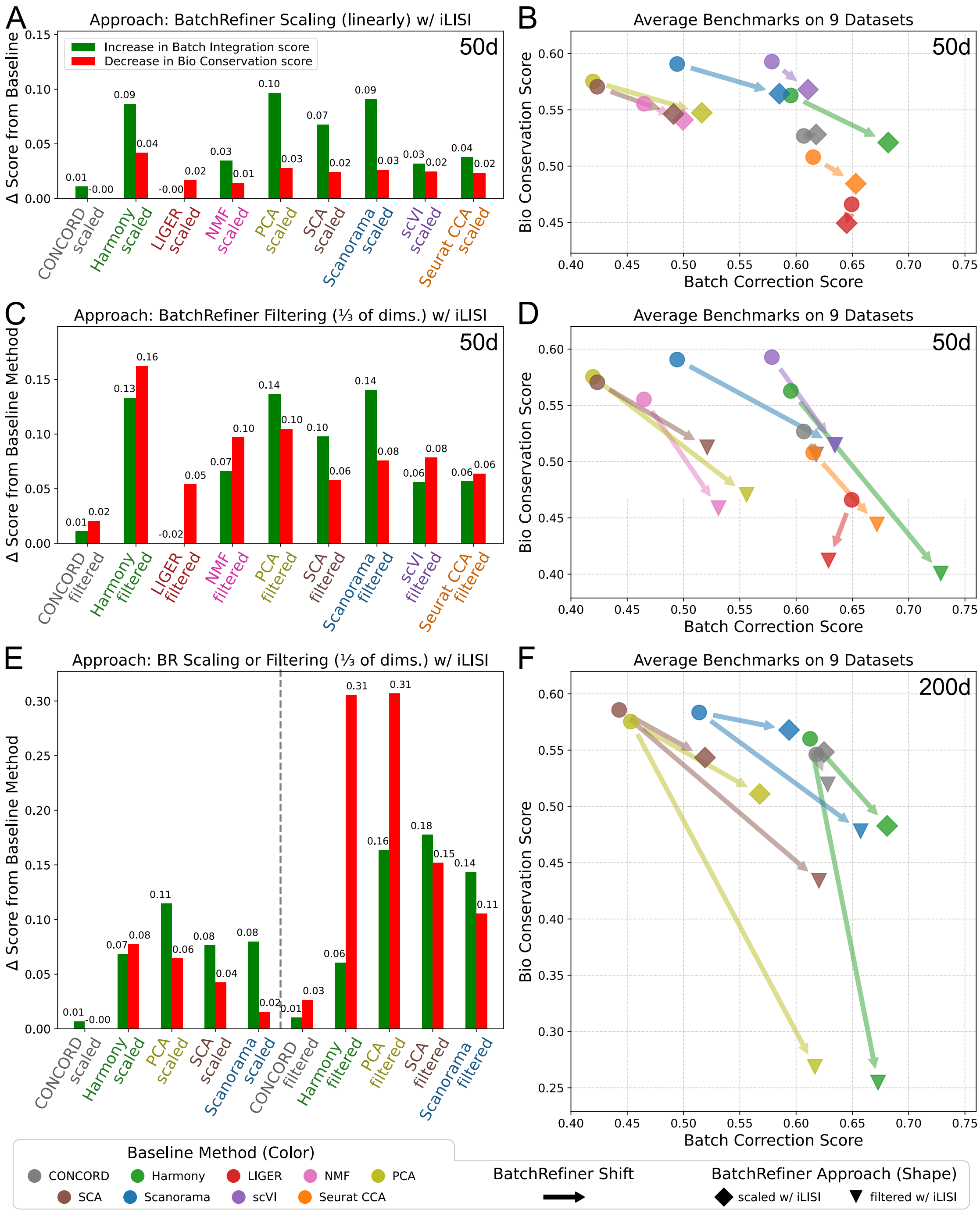


**Supplementary Fig. S17.** Benchmarking BatchRefiner with iLISI and varied initial embedding dimension. **(A)** Shifts in average batch-integration score (increases, green) and average bio-conservation score (decreases, red) relative to the corresponding baseline method for BatchRefiner **scaling** of **50-dimension** embeddings using **iLISI**. Shifts are averaged across nine scRNA-seq datasets. **(B)** Benchmarking of baseline and BatchRefiner-scaled 50d embeddings of OpenProblems datasets, shown as mean bio-conservation (y) and batch-integration (x) scores, averaged across datasets. BatchRefiner-scaled embeddings are shown as squares. Baseline 50d embeddings from nine methods are shown as circles, colored by method (bottom left). Shifts from baseline methods to the corresponding scaled embedding are shown with arrows. **(C-D)** Benchmarking BatchRefiner **filtering**, removing the lowest-scoring third of dimensions using iLISI. Data are shown as in (A-B), except that filtered embeddings are shown as in triangles. **(E-F)** Benchmarking BatchRefiner **scaling and filtering** (one third of dimensions), using iLISI, of **200-dimension** embeddings from five baseline methods. Data are shown as in (A-D).

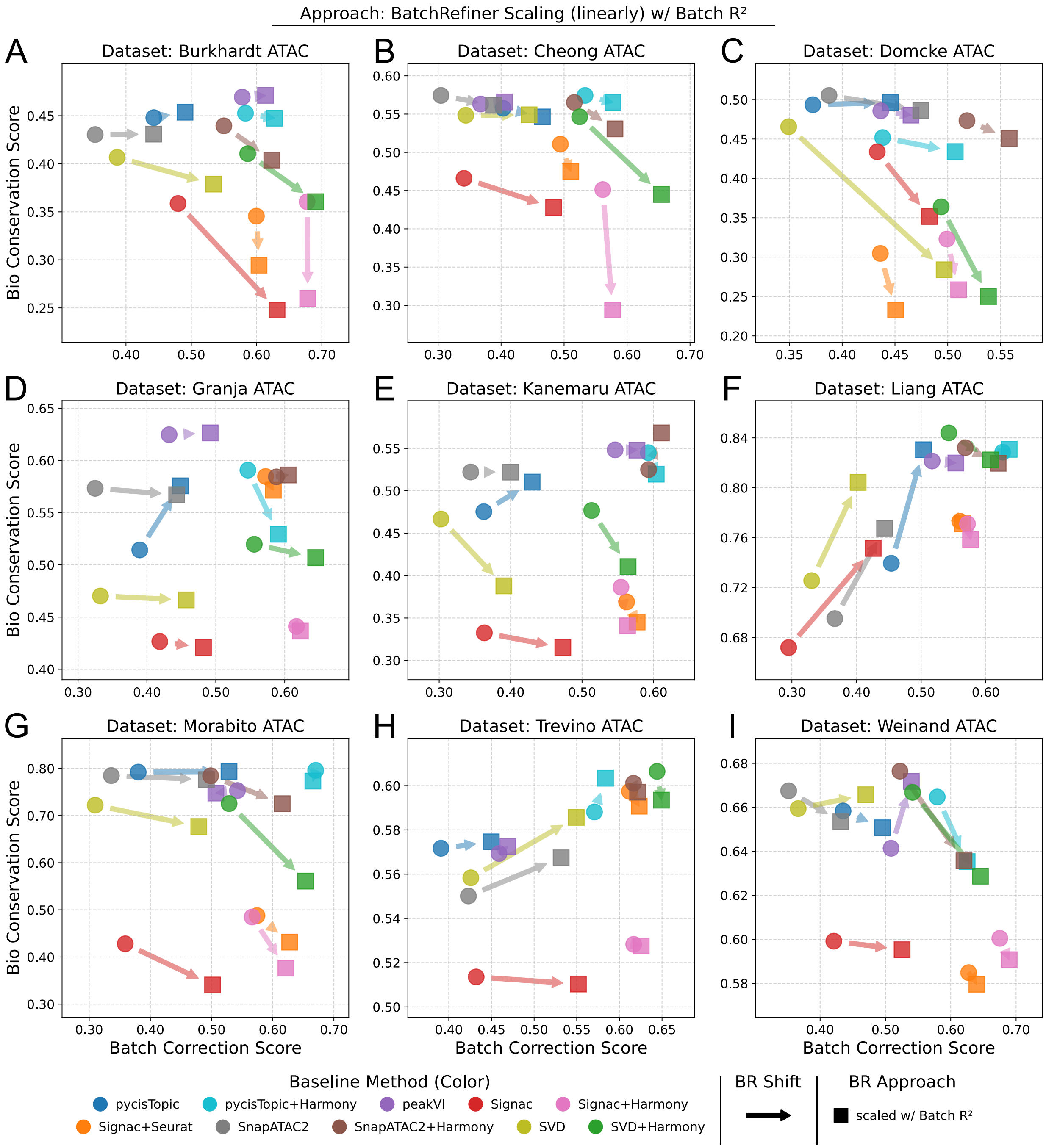


**Supplementary Fig. S18.** Benchmarks of BatchRefiner scaled embeddings on individual **scATAC-seq** datasets. Benchmarking of baseline and BatchRefiner-**scaled** embeddings, shown as mean bio-conservation (y) and batch-integration (x) scores for each of nine scATAC-seq datasets individually. BatchRefiner-scaled embeddings using batch $R^2$ as the dimension-evaluation metric are shown as squares. Baseline 100-dimensional embeddings from ten methods are shown as circles, colored by method as shown in the bottom left. Shifts from baseline methods to the corresponding scaled embedding are shown with arrows.

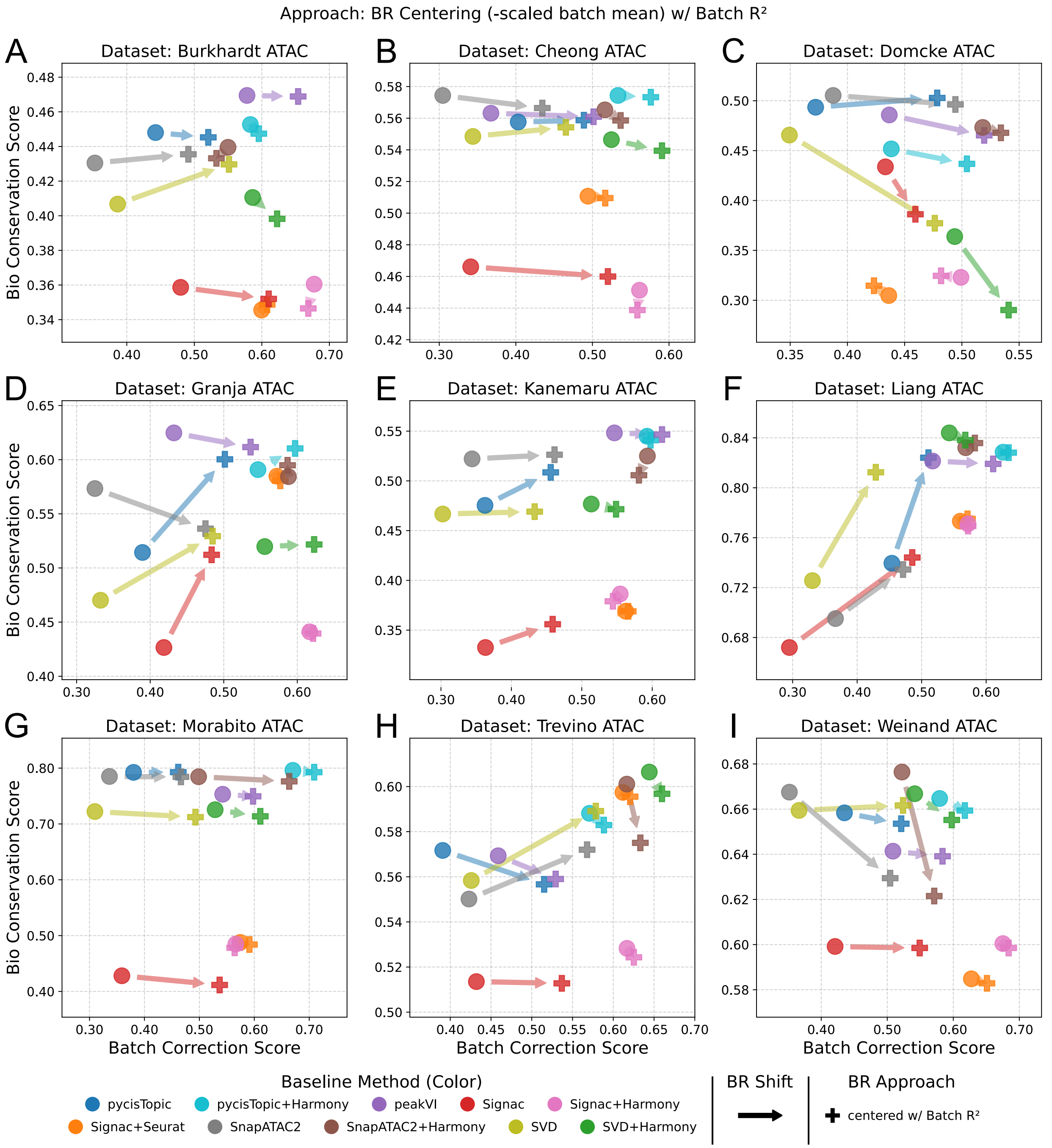


**Supplementary Fig. S19.** Benchmarks of BatchRefiner centered embeddings on individual **scATAC-seq** datasets. Benchmarking of baseline and BatchRefiner-**centered** embeddings, shown as mean bio-conservation (y) and batch-integration (x) scores for each of nine scATAC-seq datasets individually. BatchRefiner-centered embeddings using batch $R^2$ as the dimension-evaluation metric are shown as plus symbols. Baseline 100-dimensional embeddings from ten methods are shown as circles, colored by method as shown in the bottom left. Shifts from baseline methods to the corresponding centered embedding are shown with arrows.

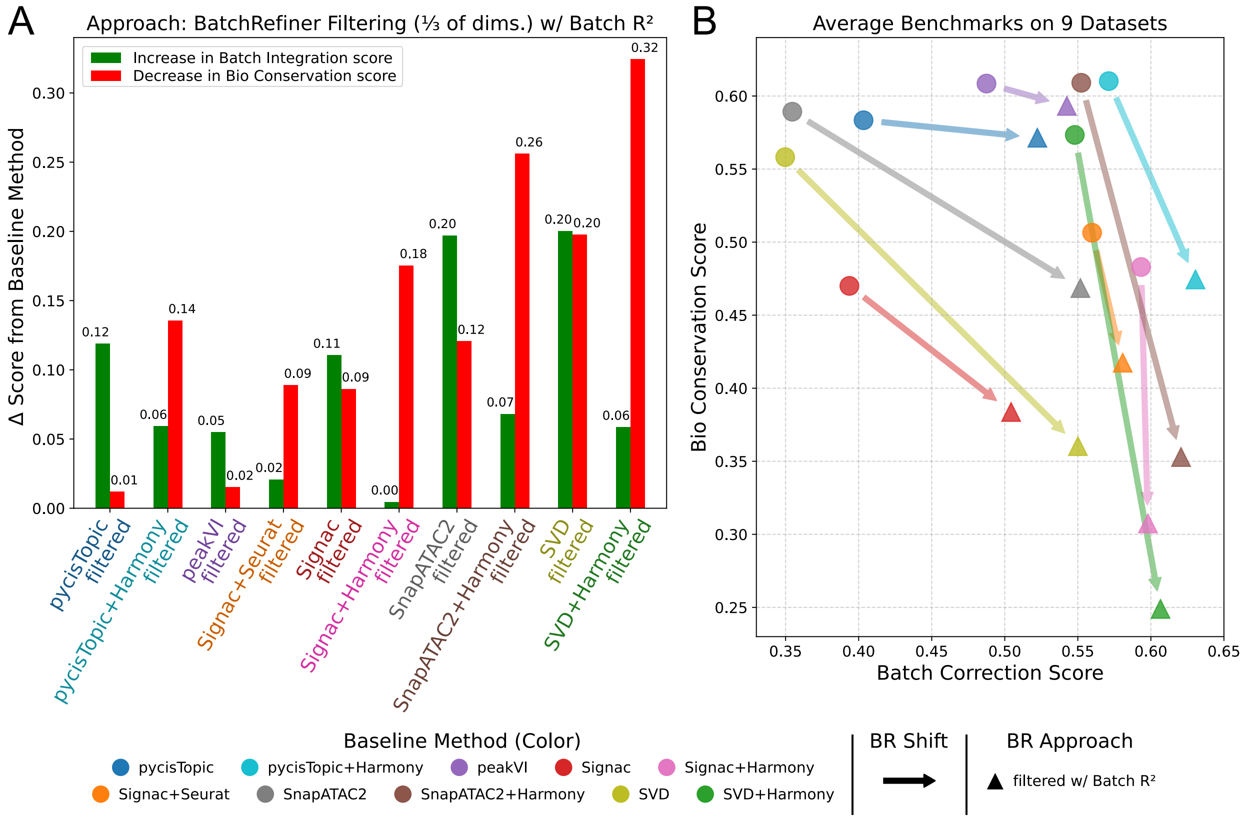


**Supplementary Fig. S20.** Benchmarking a representitive BatchRefiner **filtering** on **scATAC-seq**. **(A)** Shifts in average batch-integration score (increases, green) and average bio-conservation score (decreases, red) relative to the corresponding baseline method for BatchRefiner filtering using batch $R^2$ to remove the lowest-scoring **third** of embedding dimensions. Shifts are averaged across nine scATAC-seq datasets. **(B)** Mean bio-conservation (y) and batch integration (x) scores of baseline and BatchRefiner-filtered embeddings. BatchRefiner-filtered embeddings as in (A) are shown as triangles. Baseline 100-dimensional embeddings from ten methods are shown as circles, colored by method as shown in the bottom left. Shifts from baseline methods to the corresponding filtered embedding are shown with arrows.

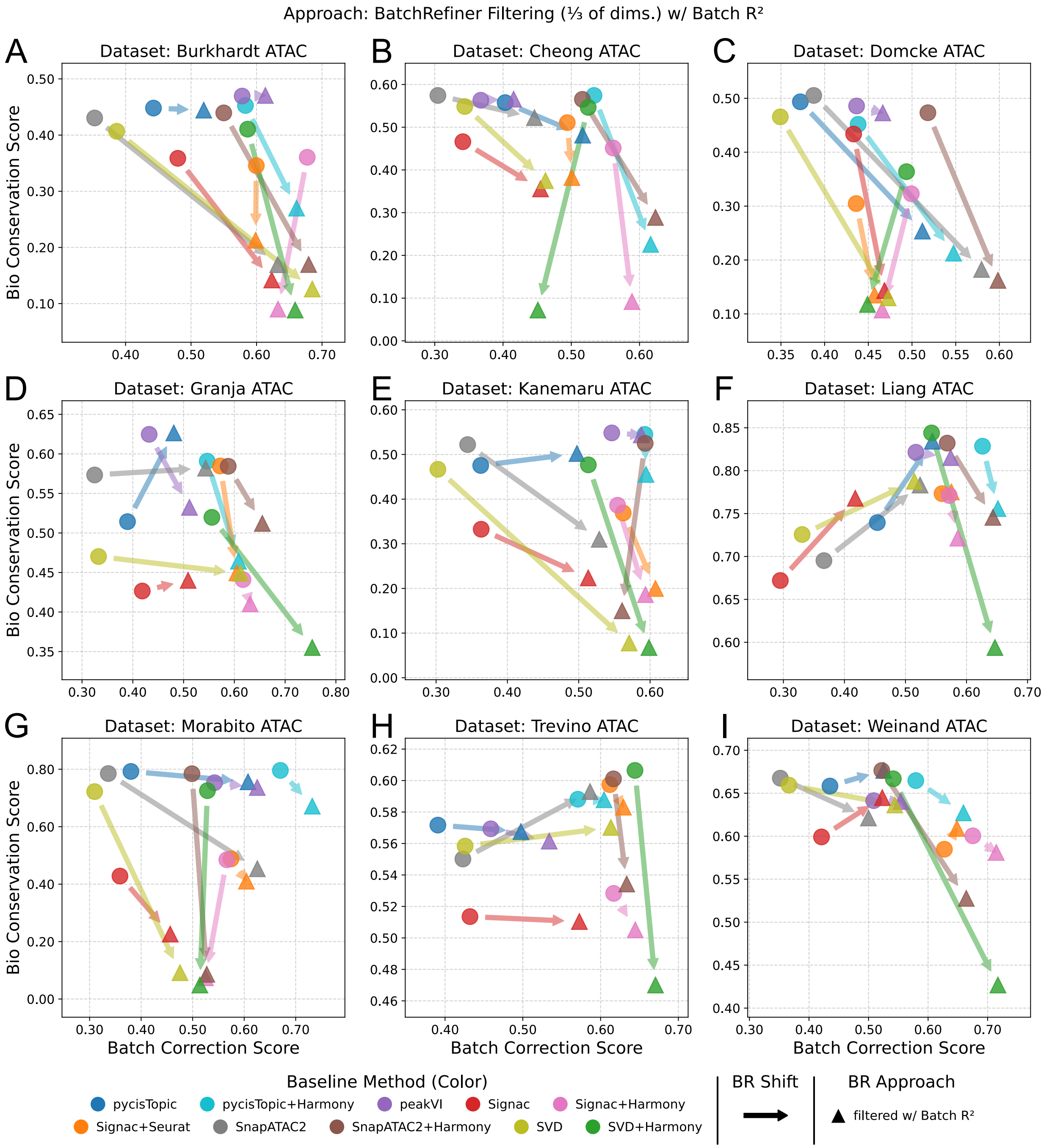


**Supplementary Fig. S21.** Benchmarking BatchRefiner filtering one third of dimensions on individual **scATAC-seq** datasets. Benchmarking of baseline and BatchRefiner-**filtered** embeddings, shown as mean bio-conservation (y) and batch-integration (x) scores for each of nine scATAC-seq datasets individually. BatchRefiner-filtered embeddings using batch $R^2$ as the dimension-evaluation metric, to filter one **third** of dimensions, are shown as triangles. Baseline 100-dimensional embeddings from ten methods are shown as circles, colored by method as shown in the bottom left. Shifts from baseline methods to the corresponding filtered embedding are shown with arrows.
